# Splice-Aware Optimization Prevents Pervasive Missplicing of Natural and Synthetic cDNAs

**DOI:** 10.64898/2026.05.25.727620

**Authors:** Mohd Ahmad, Fernando Bellido Molias, Laura Cano Aroca, Christine Mordstein, Samir Watson, Nabid Bhuiyan, Marguerite J Clarke, Chava Kimchi-Sarfaty, Upendra Katneni, Eleanor Gaunt, Laurence D Hurst, Nikolai A Netuschil, Thomas Hofmeister, Michael Liss, Grzegorz Kudla

## Abstract

Heterologous gene expression is widely used across biology and medicine, and often relies on codon optimization to increase protein yields. Here we uncover missplicing as a common and largely unrecognized failure mode of heterologous expression. Using systematically designed libraries comprising over 5,000 synthetic reporter genes and natural human cDNAs, we find that the majority of gene variants expressed in a human cell line are at least partially spliced, and in many variants the spliced isoform dominates, reducing protein output or ablating expression entirely. By analysing sequence determinants of expression across multiple human cell lines, we uncover a hierarchical architecture of regulatory control, where GC content establishes baseline mRNA levels, local sequence features influence splicing, and tissue-specific codon adaptation to tRNA pools fine-tunes translation efficiency. These findings enable us to develop predictive models of expression and splicing, benchmark current optimization strategies, and design a splice-aware optimization algorithm that substantially improves transgene performance.

## INTRODUCTION

In the genetic code, 18 out of 20 amino acids are each encoded by multiple codons, known as synonymous codons. Although synonymous codons specify the same amino acid, they are not functionally equivalent, and their usage can influence gene regulation at multiple levels. This enables "gene optimization", where codons in a gene are substituted to enhance protein expression—the most common goal—as well as other outcomes such as manufacturability, immunogenicity, and cellular burden. Gene optimization can increase the expression of target genes by 100-fold or more, making it essential in areas such as vaccine design and biomanufacturing (Gustafsson, Govindarajan, and Minshull). Optimization is typically achieved by modulating global and local coding sequence properties, such as codon usage frequencies, GC content, predicted mRNA structure, and the presence of sequence motifs, while preserving the original amino acid sequence of the encoded protein. Despite extensive research, the rules for gene optimization remain elusive (Ranaghan et al.). This is due in part to the large number of sequence features that can potentially be optimized, their unpredictable interactions with each other, and the context-dependent nature of optimization—where improvements for one outcome may lead to suboptimal performance in another. As a result, there is a pressing need for understanding the rules underlying gene optimization.

Historically, gene optimization has focused on matching codon usage patterns to cellular tRNA populations. Several metrics quantify codon usage, including the Codon Adaptation Index (CAI) (Sharp and Li), the tRNA Adaptation Index (tAI) (dos Reis, Savva, and Wernisch), and the Codon Stability Coefficient (CSC) (Presnyak et al.). These measures, while distinct, typically correlate with one another. Beyond codon usage, GC content—the proportion of guanine and cytosine nucleotides in a sequence—plays a critical role in various steps of gene expression, including transcription and transport of RNA (Zuckerman et al. ; Palazzo and Kang)(Courel et al.). GC content also influences the thermodynamic stability of mRNA, which can affect both translation efficiency and RNA turnover (Kudla et al. ; Zhang et al.). The frequency of CpG dinucleotides is well known for its role in transcriptional regulation, but also affects mRNA stability through interactions with RNA-binding proteins, such as ZAP (Takata et al.). Importantly, these parameters are intrinsically linked: modifications to one invariably affect the others. For instance, CpG content correlates with codon pair bias (Tulloch et al. ; Kunec and Osterrieder), while codon usage patterns are inherently connected to nucleotide composition, RNA structure, and exonic splice enhancers (ESEs) (Parmley, Chamary, and Hurst), creating a web of relationships that must be considered during optimization.

While most optimization efforts have focused on enhancing translation, synonymous recoding can lead to unwanted consequences, such as changes in protein folding, immunogenicity, or posttranslational modifications (Katneni et al.). Synonymous substitutions can also inadvertently introduce cryptic splice sites, disrupt splicing enhancers and silencers, or alter RNA secondary structures, potentially leading to exon skipping or creation of short *de novo* introns, known as exitrons (Marquez et al.). Numerous studies have shown that single-nucleotide changes in endogenous human genes can induce splicing defects with clinical consequences (Radrizzani et al. ; Zhang et al.). Similar splicing disruptions have been documented in recombinant constructs, including epitope-tagged reporters and codon-optimized SARS-CoV-2 and HIV genes (Ficarelli et al. ; Kowarz et al. ; Ansseau et al. ; Paget-Bailly et al.). Despite this, gene optimization pipelines rarely incorporate splicing prediction models, and in laboratory practice, codon-optimized genes are not commonly assessed for the presence of missplicing. More worryingly, the possibility that codon optimization might introduce unwanted protein forms is not routinely considered in clinical applications, where transgenes are introduced into patients’ DNA for therapeutic purposes.

Here, using large libraries of reporter variants, we dissect the individual and combined effects of codon usage, nucleotide composition, and local sequence motifs on transcript abundance, integrity, and translation efficiency in human cells. Our results reveal unintended splicing as a pervasive cause of expression loss and provide a framework for splice-aware gene optimization that improves both yield and transcript fidelity.

## RESULTS

### Design and Validation of a Systematic Codon Analysis Platform

To systematically evaluate how synonymous codon usage influences gene expression in human cells, we developed a modular platform that integrates large-scale variant libraries with orthogonal readouts of mRNA abundance, stability, splicing, and translation efficiency (**Fig. 1**). The platform comprises multiple components: (1) an arrayed library of 194 factorially designed GFP variants (GFP_DOE) that explores the individual and combined effects of GC content, CpG dinucleotide frequency, and codon adaptation index (CAI); (2) a pooled library of 4,675 synthetic GFP variants with high GC and CAI (GFP_optimized); (3) smaller libraries of mKate2 variants and therapeutic transgenes used in genetic medicine; and (4) a pool of 689 natural human coding sequences from the human ORFeome. These libraries were cloned into barcoded expression plasmids, enabling transfections into multiple human cell lines and unbiased quantification of RNA expression and splicing via long-read sequencing, and protein expression of GFP variants via fluorescence measurements.

**Figure 1.**
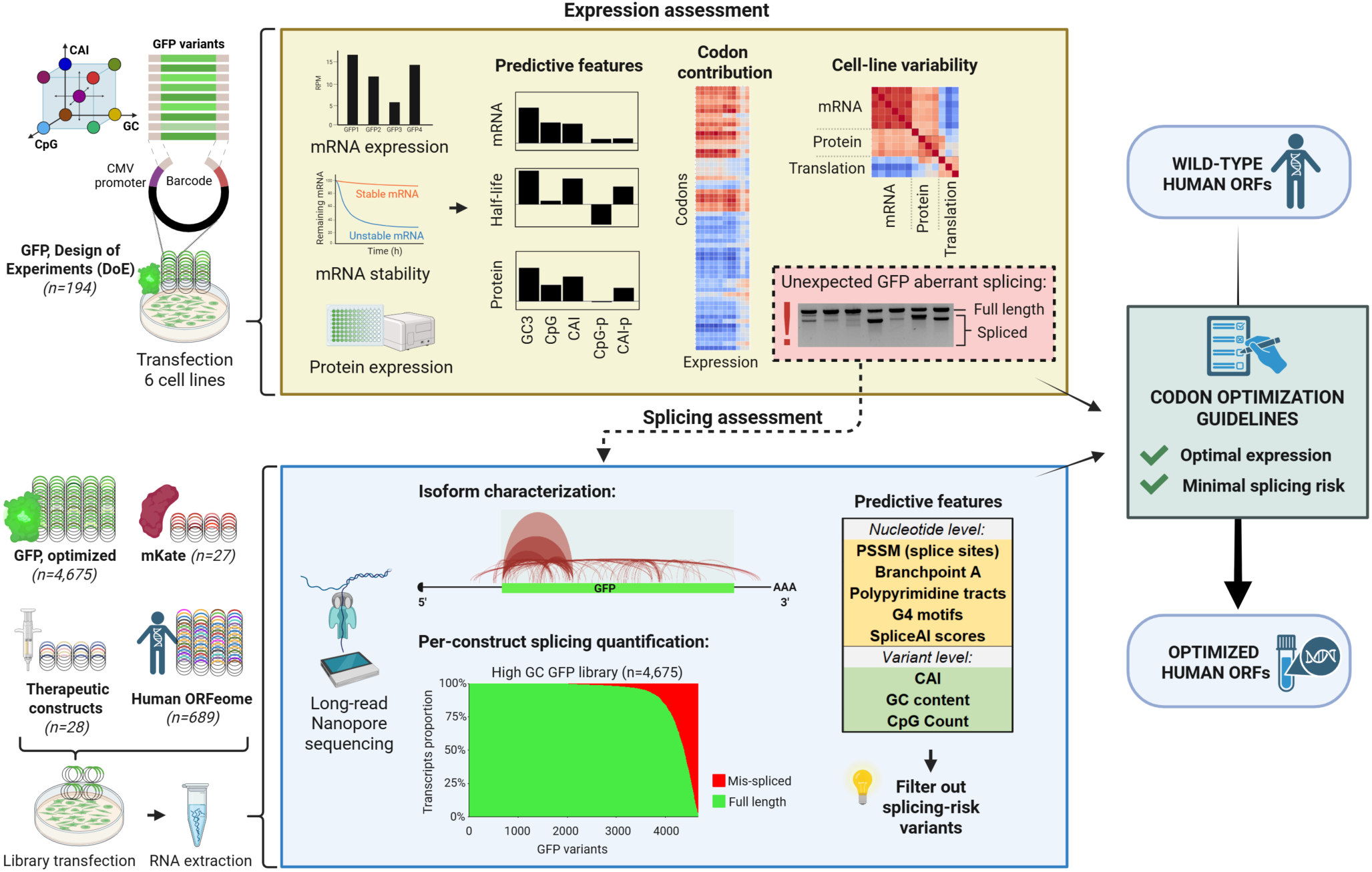
Design of systematic codon analysis platform.

We initially employed the GFP_DOE library to analyse the sequence determinants of mRNA and protein expression in HeLa cells. The library employed a factorial design of experiments (DOE) strategy, where GC, CpG and CAI were each tested at multiple levels to establish causal effects on expression (**Fig. 1**, **Fig. S1A**). The variants were designed to span the natural variation observed in human genes (**Fig. S1B**) and to maximize sequence divergence, resulting in an average of 147 nucleotide substitutions between pairs of variants. This design allowed us to disentangle the effects of highly correlated sequence features and assess their individual contributions to gene regulation (**Fig. S1C**). To ensure experimental robustness, each variant was synthesized individually, barcoded, and pooled in equimolar amounts prior to transfection (**Fig. S1D**). Equal representation was confirmed by long-read DNA sequencing and results were cross-validated across technical replicates and independent quantification methods (**Fig. S2A, B, C**).

### Dissecting the Causal Relationships Between Sequence Features and Gene Expression

Initial experiments in HeLa cells revealed large variation in mRNA and protein expression across the library (**Fig. 2A, B, S2D, E**). We observed consistent grouping based on parameter values, with variants sharing similar GC, CAI, and CpG levels displaying comparable expression profiles. Similar results were obtained in a Flow-Seq assay in which the variants were stably integrated into a genomic landing pad in HeLa cells (**Fig. S2F**), indicating that genomic context and mode of transfection does not affect the conclusions, similar to previous studies (Kudla et al. ; Mordstein et al.). The complete sequences and characterization data for all variants are provided in **Supplementary Dataset 1**.

**Figure 2.**
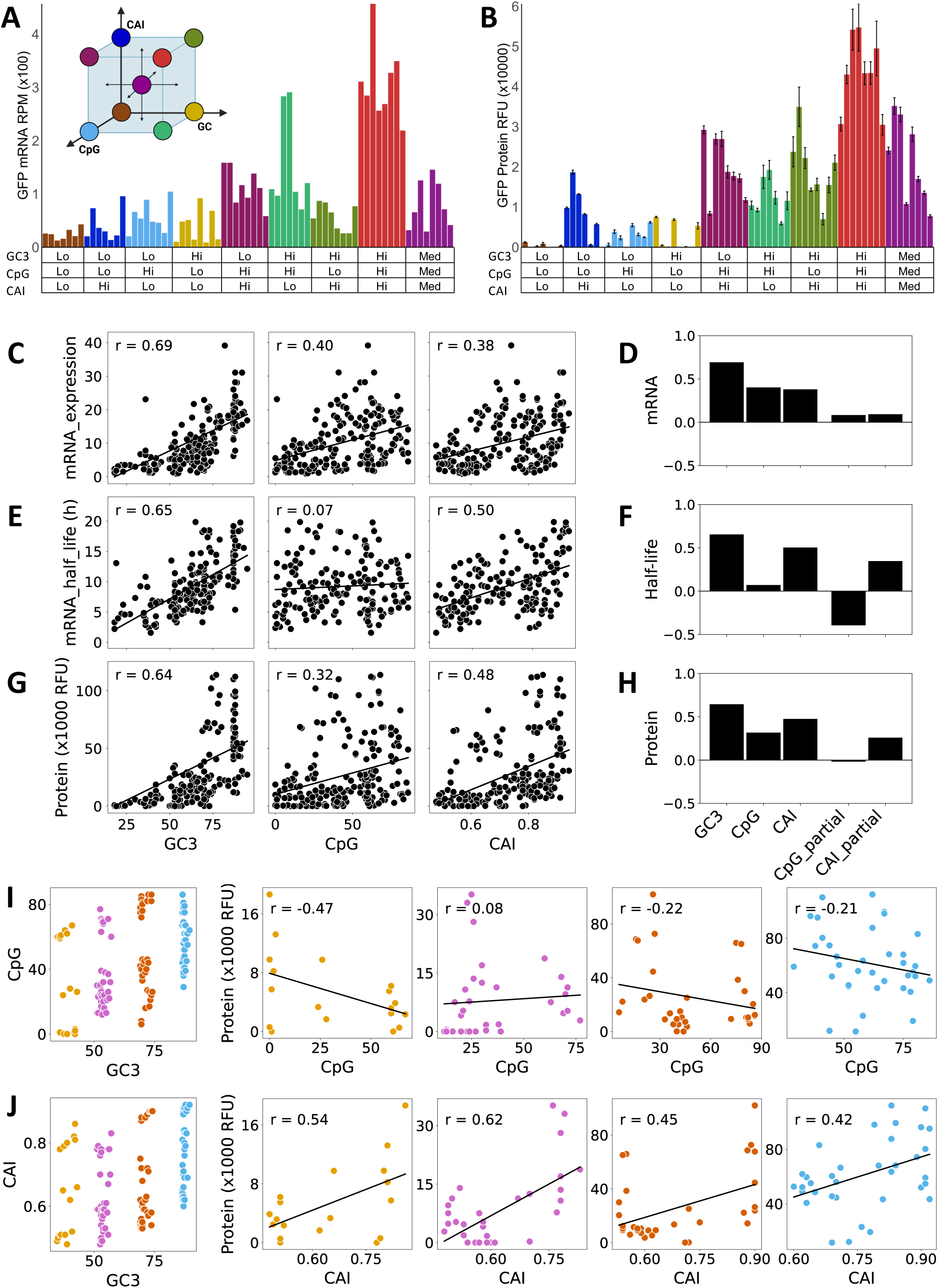
Sequence determinants of GFP expression in HeLa cells. **A)** Transient mRNA expression of 72 GFP variants in HeLa cells (RPM = reads per million). Inset, distribution of variants in a 3D parameter space. **B)** Transient protein expression of the same variants (RFU = relative fluorescence units). GC3, CpG, and CAI have low (Lo), medium (Med), and high (Hi) levels. **C, E, G)** Correlations of mRNA expression, mRNA half-lives, and protein expression of 194 GFP_DOE library variants with sequence features. **D, F, H)** Pearson correlations and partial correlations controlling for GC content of sequence features with expression levels. **I, J)** Stratifying variants according to their GC3 content reveals a negative correlation of protein expression with CpG (I) and positive correlation with CAI (J).

We next analysed the impact of global sequence properties on expression. GC content showed the strongest correlation with mRNA levels (R=0.69), while CpG content and CAI showed moderate correlations (R= 0.40 and R= 0.38 respectively, **Fig. 2C**). Partial correlation analysis, controlling for GC content, revealed that CAI and CpG had, at best, small independent effects on mRNA levels (**Fig. 2D**). This conclusion was further supported by analysing expression variation within GC-content-matched subgroups of variants (**Fig. S3 A, B**). Notably, the relationship between GC content and expression was nonlinear: variants with GC3<0.5 typically showed low expression, while those with GC3>0.5 exhibited variable mRNA levels, suggesting that high GC3 content is necessary but not sufficient for efficient mRNA production.

To determine if these patterns arise from altered RNA synthesis or degradation, we performed RNA stability measurements. We transfected HeLa cells with pooled GFP variants, treated cells with the transcription inhibitor triptolide, and quantified mRNA levels over time using direct cDNA sequencing. Triptolide was chosen over actinomycin D as it lacks bias against GC-rich sequences. Analysis of control transcripts confirmed expected half-lives for actin and GAPDH (>8h) and c-myc (∼1h) (**Fig. S3 C**). GFP mRNA half-lives correlated strongly with steady-state RNA levels and, like these, were primarily determined by GC content. After controlling for GC content, CAI showed a slight positive effect on stability while CpG content showed a negative effect (**Fig. 2E, F, S3 D,E,F**). At the protein level, measurements revealed strong correlation with GC content (R=0.65), followed by CAI (R=0.48), and weaker correlation with CpG (R=0.32) (**Fig. 2G**). Partial correlation analysis showed that both GC content and CAI independently influenced protein levels (**Fig. 2H**). Examination of GC-content-matched subgroups confirmed these relationships, demonstrating that CAI effects on protein levels persist when controlling for GC content, while CpG had no effect or a negative effect (**Fig 2I, J**).

**Figure 3.**
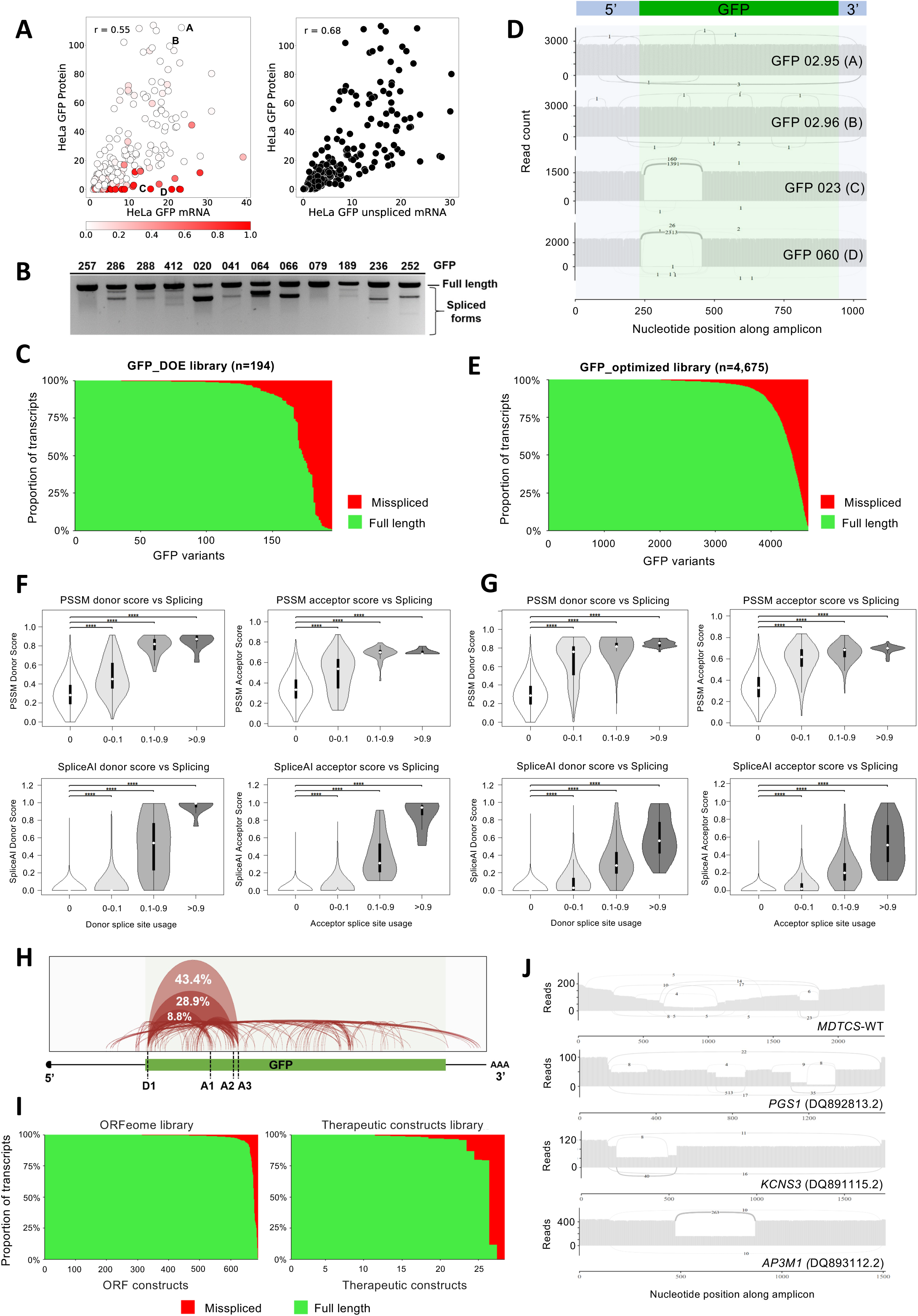
Heterologous cDNAs are frequently misspliced. **A)** (Left) Correlation between GFP protein and total GFP mRNA expression in HeLa cells. Red colour intensity indicates the proportion of spliced GFP transcript. Representative examples of low (A–B) and high (C–D) splicing are highlighted. (Right) Correlation between GFP protein abundance and estimated levels of full-length GFP mRNA. **B)** RT-PCR of RNA from selected GFP variants showing lower bands consistent with the presence of splicing. Numbers on top indicate IDs of variants from the GFP_DOE library and from a previous study (Mordstein et al., 2020). **C)** Proportion of spliced GFP transcripts across the GFP_DOE library. **D)** Sashimi plots of highlighted variants from (A), confirming splicing at read level. **E)** Proportion of spliced transcripts across the GFP_optimized library. **F)** Violin plots of donor and acceptor PSSM (top) and SpliceAI (bottom) scores across all GFP variants and positions in the GFP_DOE library, stratified by splice site usage. Significance was tested with Wilcoxon rank-sum tests (****p < 0.0001). **G)** Violin plots of PSSM and SpliceAI scores as in (F), for the GFP_optimized library. **H)** Diversity of splicing events in the GFP_optimized library. Arc thickness reflects frequency. **I)** Proportion of spliced transcripts arising from transfection of human cDNA vector libraries: ORFeome (left) and therapeutic cDNAs (right). **J)** Sashimi plots demonstrating prevalent splicing in transcripts expressed from wild-type cDNAs in the therapeutic collection (*MDTCS*-WT, top) and the ORFeome set (*PGS1*, *KCNS3*, *AP3M1*, bottom).

### Heterologous Genes are Frequently Misspliced

Although GFP protein and mRNA levels correlated with each other, we noticed a subset of variants that showed low fluorescence despite moderate or high mRNA levels (**Fig. 3A, left**). Analysis of the corresponding cDNAs by RT-PCR and agarose gel electrophoresis revealed lower bands consistent with the presence of spliced products (**Fig. 3B**). To evaluate splicing, we performed pooled transfection of the library into HeLa cells followed by PCR and long-read Nanopore sequencing. We found evidence of missplicing (>0.1%) in 184 out of 194 variants in the library, including 57 variants with more than 5% spliced reads, and 22 variants where the majority of reads were spliced (**Fig. 3C**). Splicing was confirmed through examination of individual reads (**Fig. 3D**) and the detection of canonical splice sites flanking the missing region of the cDNA (**Fig. S4**). GFP protein abundance correlated better with estimated full-length mRNA levels (total mRNA reads × fraction unspliced; r=0.68) than with uncorrected total mRNA reads (r=0.55) (**Fig. 3A, right**), confirming that missplicing explains part of the protein–RNA discrepancy.

We then investigated whether known sequence motifs predict splicing outcomes. Using Position-Specific Scoring Matrices (PSSMs) derived from human mRNAs, we found that splice donor and acceptor PSSM scores were strong predictors of splicing activity and correlated significantly with measured splicing events across all GFP variants (p < 0.001) (**Fig. 3F**). This was also true when the analysis was limited to positions compatible with canonical splicing (i.e. positions with GT and AG dinucleotides for donor and acceptor sites, respectively) (**Fig. S5**).

Similarly, PSSM scores for branchpoint motifs, polypyrimidine tracts, and G-quadruplex (G4) also correlated with the splicing usage of nearby cryptic splice sites (**Fig. S5-S7**), underscoring their essential role in enabling efficient splice site recognition. In addition, we observed a higher frequency of exonic splicing enhancers (ESEs) in GFP constructs showing moderate to high levels of splicing (>10%) (**Fig. S8**). Features such as GC content, CAI, and CpG frequency were statistically associated with splicing, though their effect sizes were minimal (**Fig. S9**). We next turned to spliceAI, a machine learning tool trained to predict splicing patterns in endogenous genes. We observed a strong correlation of the spliceAI-predicted donor and acceptor scores with observed splicing efficiency across the library (**Fig. 3F**), suggesting that, although spliceAI has not been trained on synthetic transgenes, it can nevertheless identify splicing patterns in this context.

To test if these findings generalize to codon optimized genes, we used the pooled GFP_optimized library, featuring an average GC3 content of 84% and average CAI of 0.75, as these values typically resulted in high protein yields in our experiments (**Fig. 2B**). Using RT-PCR and Nanopore sequencing, we obtained a coverage of ≥20 reads per variant for 4,675 variants. This revealed a profile of spliced and unspliced cDNAs similar to the GFP_DOE library (**Fig. 3E**). We identified hundreds of distinct splice isoforms, with a few isoforms explaining the majority of splicing activity **(Fig. 3H**). With the larger size and particular design of this GFP collection, we confirmed that splicing strongly correlated with PSSM-derived and spliceAI-derived donor and acceptor scores (**Fig. 3G**), as well as polypyrimidine tracts, branchpoint A motifs, G-quadruplexes, and to some degree with GC content, CAI, and CpG frequency (**Fig. S10-S13**).

To evaluate if unintended splicing is a general feature of transgene expression, we analysed a broad set of constructs beyond GFP, including 27 mKate variants, 28 cDNAs of therapeutic proteins (wild-type and codon-optimized *ADAMTS13*, *MDTCS*, and coagulation factors VIII and IX), and 689 intronless human cDNAs from the ORFeome collection. Splicing was consistently detected across all datasets (**Fig. 3I, S14**): in the mKate library, all constructs exhibited detectable splicing, and 11.1% (3/27) displayed >5% spliced reads (**Fig S14**). Among the therapeutic constructs, 82.1% (23/28) exhibited detectable splicing, with the highest levels observed in wild-type ADAMTS13 and MDTCS (**Fig. 2I, J**). Despite being intronless and derived from already spliced mRNAs, 6.7% of the ORFeome entries (46/689) showed >5% splicing, and 2% (14/689) presented >50% splicing penetrance (**Fig. 2I, J**). Mechanistically, some spliced ORF transcripts matched annotated isoform junctions, particularly when long ORF exons overlapped regions corresponding to exon–intron–exon structures in a major gene isoform **(Fig. S15A, B).** Yet, most aberrant splicing events in the ORFeome library originated from cryptic splice site activation following intron removal **(Fig. 3J, S15C, D)**. Taken together, these results show pervasive splicing of cDNAs expressed in a human cell model, and that unintended splicing is not confined to synthetic or codon-optimized constructs, but also affects native human cDNAs. The associations between motif-derived scores and splice site usage open the possibility of integrating splicing-aware filters into codon optimization pipelines.

### Conservation of Sequence Effects Across Human Cell Lines

To investigate if these results are consistent across cell types, we measured mRNA expression levels of all GFP variants in five additional human cell lines: HEK293 (embryonic kidney), SH-SY5Y (neuroblastoma), A549 (lung epithelial), HepG2 (hepatocellular carcinoma), and RPE (retinal pigment epithelium). Despite their distinct tissues of origin, GFP mRNA expression profiles were remarkably similar, with pairwise correlations across cell lines consistently exceeding 0.9 (**Fig. 4A**). As observed in HeLa cells, GC content showed the strongest correlation with mRNA expression in all cell lines tested, while CAI maintained a modest correlation after controlling for GC content (**Fig. 4B**).

**Figure 4.**
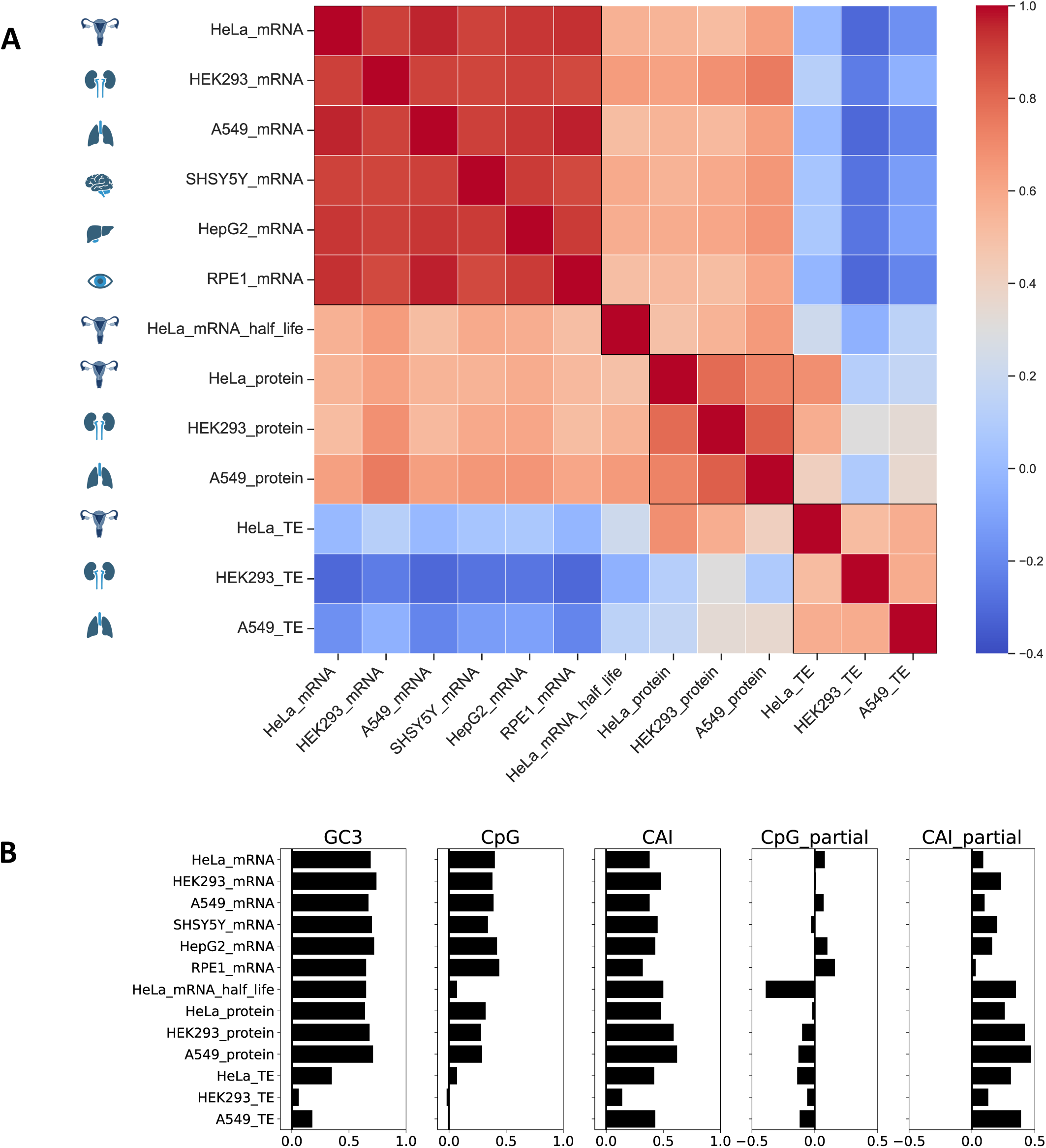
Sequence determinants of GFP expression across cell types. **A)** Correlation matrix of GFP mRNA expression, half-life, protein expression, and translation efficiency among the six cell lines tested. **B)** Pearson correlations and partial correlations (controlling for GC content) of GFP gene features with expression levels.

Cell-type specific effects became more apparent when analysing protein levels **(Fig. 4A)**. While correlated with mRNA levels, protein expression showed lower correlation between cell lines than mRNA did. We hypothesized that this is due to tissue-specific codon preferences that may influence translation rates. To test this, we analysed translation efficiency (TE), defined as the protein-to-mRNA ratio. Indeed, this revealed more distinct patterns, with varying effects of predictive features on translation (**Fig. 4B**). Despite these differences, CAI was the most predictive feature for TE in all cell lines tested, an effect that persisted after accounting for other parameters. In contrast, GC3 showed a weaker association with TE that largely disappeared after controlling for CAI (**Fig. S16A**). As seen in HeLa cells, GFP protein levels showed higher correlations with the estimated level of unspliced GFP mRNA than with total GFP mRNA in HEK293 and A549 cells, suggesting that splicing patterns are conserved across cell lines (**Fig. S16B, C**).

We also investigated how sequence determinants of expression depend on the physiological state of cells or on the presence of specific RNA-binding proteins. Following previous observations that serum starvation alters tRNA pools and induces a shift from proliferative to non-proliferative cell states (Gingold et al.), we performed serum withdrawal experiments in HeLa cells. Although serum starvation reduced overall GFP mRNA abundance, variant expression levels remained highly correlated between conditions (R>0.9, **Fig. S17 A**). We also asked whether expression of GFP variants is modulated by ZAP, an RNA-binding protein known to target CpG-rich viral transcripts. Upon expressing a subset of GFP variants in cells lacking ZAP or TRIM25, a cofactor in the ZAP RNA degradation pathway, we observed no significant effect on CpG-rich GFP variants (**Fig. S17 B-D**).

### Codon-Specific Contributions to Gene Expression

The cell-type specific patterns in translation efficiency suggested that individual codons might have distinct effects on gene expression that vary between cell types. To systematically investigate these effects, we calculated Codon Expression Coefficients (CEC), a metric that quantifies the correlation of codon frequencies with expression, inspired by the CSC score (Presnyak et al.). We calculated separate coefficients for mRNA levels (CEC_mRNA_), protein levels (CEC_protein_), and translation efficiency (CEC_TE_). Many codons showed distinct positive or negative CEC: C-ending codons generally associated with positive scores, A- and T-ending codons with negative scores, while G-ending codons showed variable effects (**Fig. 5A**). CEC_mRNA_ and CEC_protein_ were highly similar between cell lines while CEC_TE_ were more distinct. Despite differences in CEC_TE_ across cell types, protein expression was highly correlated between cells (**Fig. 4A**). This likely reflects the fact that, in this heterologous system, protein abundance is driven largely by mRNA levels. Similar patterns were observed when quantifying the correlation of codon usage on mRNA expression of ORFeome constructs (**Fig. 5B**), but correlations were weaker, presumably reflecting contributions of additional factors beyond codons bias.

**Figure 5.**
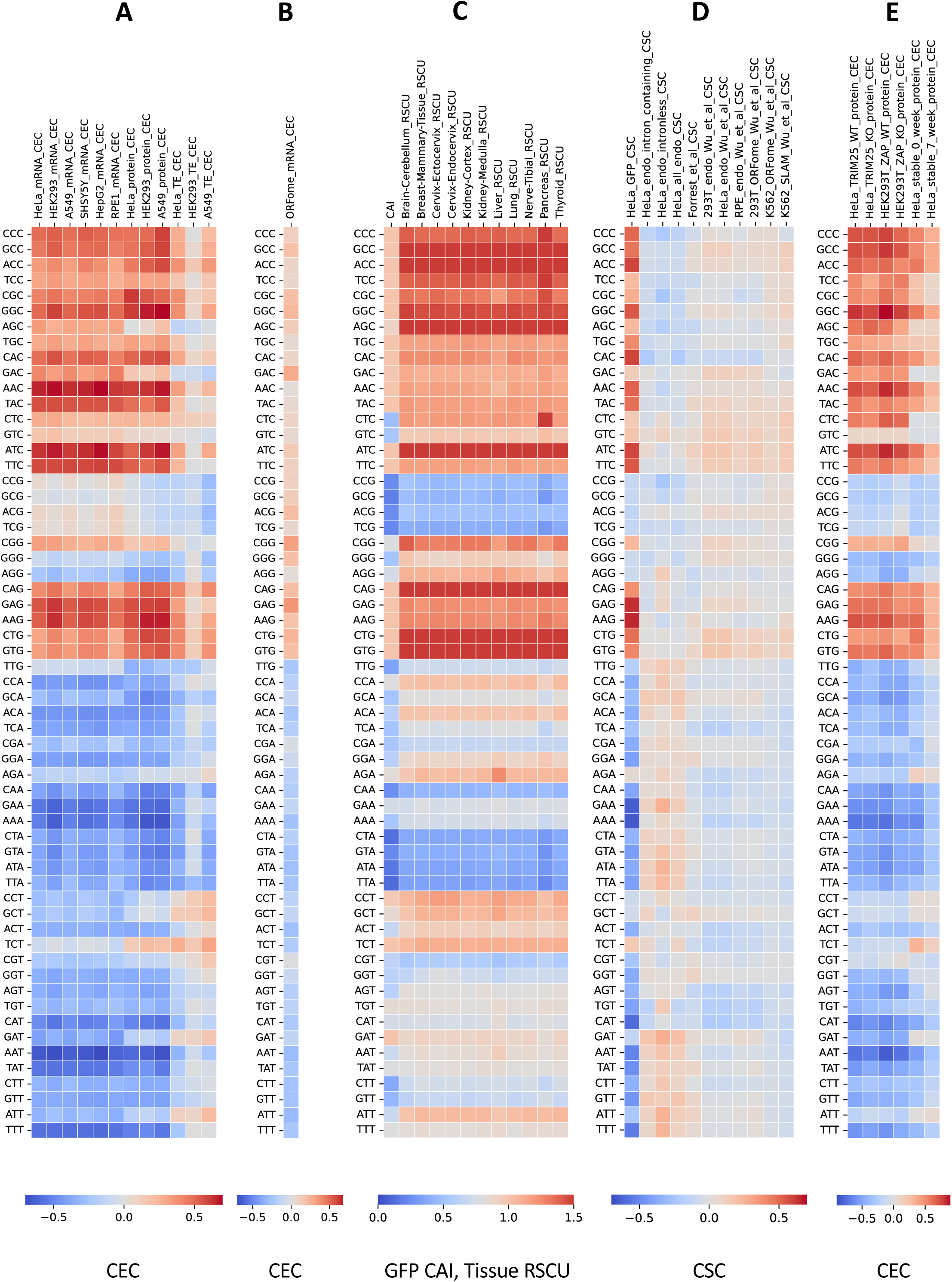
Comparison of codon expression coefficients with established codon scores. **A)** Codon Expression Coefficients for the mRNA, protein, and translation efficiencies across human cell types. **B)** Codon Expression Coefficients for the ORFeome mRNA. **C)** Codon Adaptation Index and RSCU scores across human tissues. RSCU scores were calculated by retrieving the tissue specific codon data from the TissueCoCoPUT database. **D)** Codon Stability Coefficients from the present study and other studies. **E)** GFP CEC calculated from the TRIM25 WT & KO cell lines, ZAP WT & ZAP KO cell lines, and from HeLa cells stably expressing GFP variants, at 0 week and 7^th^ week.

We next compared CEC scores with established codon-based metrics. CEC showed similar patterns to metrics derived from endogenous genes, including codon usage in highly expressed transcripts (CAI) and relative synonymous codon usage (RSCU) (**Fig. 5C**). The similarity was particularly apparent for CEC_TE_ scores: for example, the frequency of CCT, GCT, GAT and ATT codons correlated negatively with mRNA and protein abundance, but positively with translation efficiency, consistent with the relatively high usage of these T-ending codons in human mRNAs (**Fig. 5C**). CEC scores derived from our reporter assays showed a modest positive correlation with RNA stability (CSC) scores previously measured for endogenous genes, as well as a notably broader range than CSC scores (**Fig. 5D**). This suggests that RNA stability measurements in endogenous genes are not necessarily predictive of heterologous protein yields. This difference persisted even when we recalculated CSC using only intronless endogenous genes (**Fig. 5D**). CEC scores were also similar following deletion of ZAP and TRIM25, and for variants stably expressed from a genomic locus, showing the robustness of codon-dependent regulatory effects across various heterologous systems (**Fig. 5E**).

### Evaluating Gene Optimization Methods

A wide variety of gene optimization tools have been developed over the years, incorporating combinations of codon preference scores, RNA structure predictions, and other sequence-derived features, but the relative performance of these methods has not been systematically evaluated. We therefore developed a data-driven approach to benchmark gene optimization methods.

In the first step, we used our dataset to train predictive models of gene expression and splicing. We compared several model architectures and sets of predictive features, training each model on 80% of the data and evaluating its performance on the remaining 20%. Models based on a single predictive feature explained up to 46% of variance in GFP protein expression, with GC content being the strongest predictor (**Fig. 6A**). Among models with multiple predictive features, random forest (RF) models using codon frequencies as features performed best (R²=62% to 75% across three cell lines) (**Fig. 6A, S18**), while RF models based on global sequence properties—GC, CAI, CpG, and predicted mRNA folding energy—yielded R² values between 58% and 73%. These models generalized well across cell types and expression modalities: for example, a model trained on GFP protein expression in HeLa cells explained 62% of variation in protein expression and 57% variation in mRNA expression in HEK293 cells, for unseen variants (**Fig. S18B**). SpliceAI-predicted maximum donor and acceptor site scores for each variant accounted for 45% of the variation in splicing, 35–43% of variation in protein levels, and 21-41% of variation in mRNA levels (**Fig. 6A, S18A, S19**).

**Figure 6.**
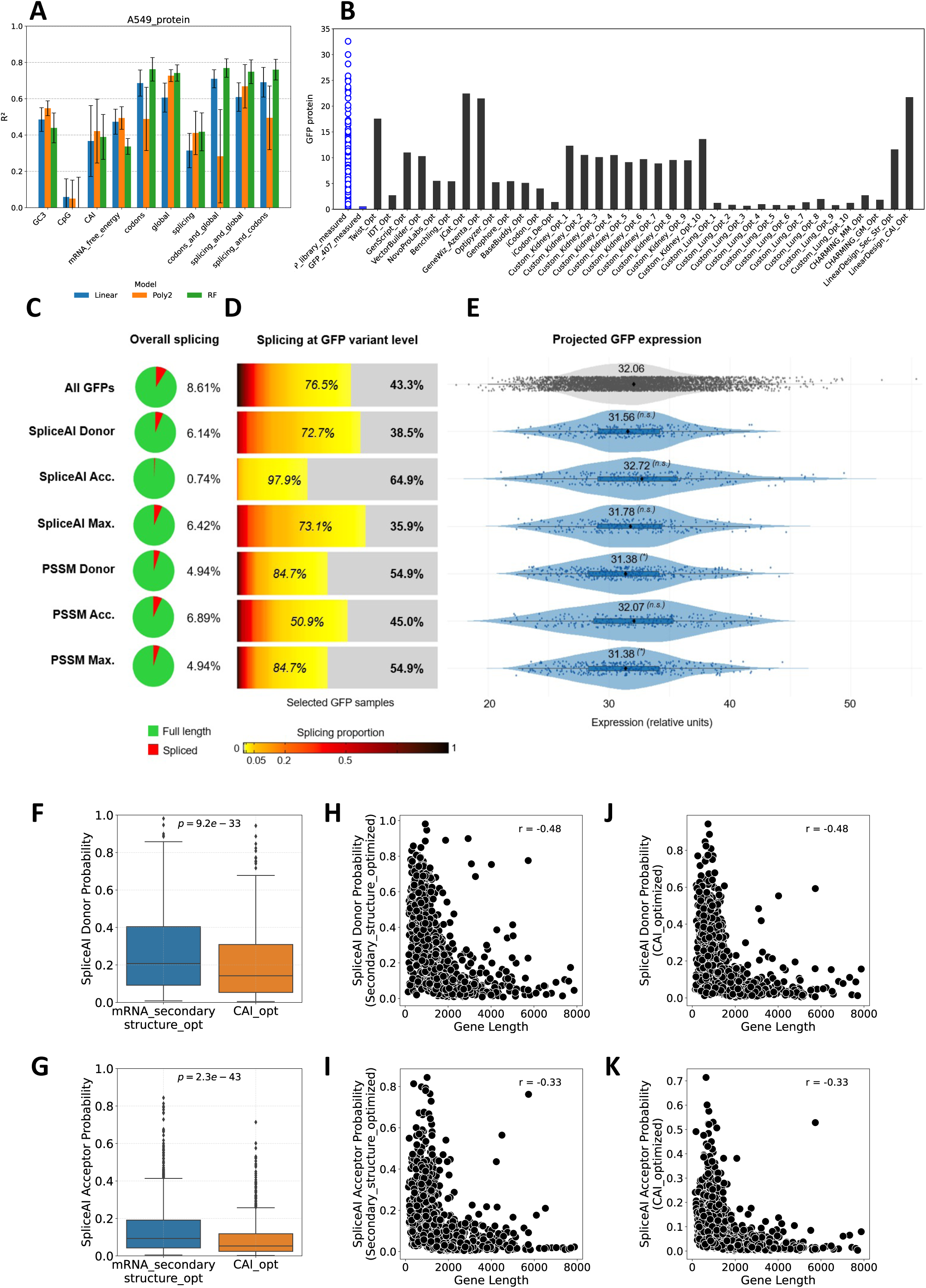
Benchmarking gene optimization methods. **A)** Out-of-fold performance of three model architectures in predicting GFP protein expression levels in A549 cells, using ten sets of predictive features. **B)** Measured protein expression in A549 cells of GFP_DOE library and GFP_407 variant (blue jitter plot and the blue bar). Predicted protein expression in A549 cells of variant GFP_407 optimized and deoptimized using several codon optimization tools, calculated using the RF_codons model (black bars). **C)** Proportion of spliced (red) versus full-length (green) transcripts across the GFP_optimized library (top) and subsets filtered by different splicing prediction criteria: 5% of variants with the lowest maximum-score SpliceAI and PSSM scores along their full sequence, for donor, acceptor, or the highest -*“max”-* of both. **D)** Distribution of GFP variants by their proportion of splicing, ranging from null (grey), low (yellow) to high (black) levels (colour scale below). The percentages on the right indicate the fraction of GFPs with no detectable splicing, while those on the left represent variants with up to 5% splicing. **E)** Predicted GFP expression levels based on the RF_codons model for the full GFP_optimized set (top, grey) and filtered subsets (blue). Statistical significance calculated with t-test: p < 0.05 (*); n.s., not significant. Acc., acceptor; Max., maximum. **F, G)** SpliceAI-predicted maximum acceptor and donor probabilities in human genes from the MANE database optimized by Linear Design. Genes were optimized to have either high mRNA secondary structure or high Codon Adaptation Index. Statistical comparisons were performed using Wilcoxon rank-sum tests. **H, I)** Correlations between gene lengths and the SpliceAI maximum donor or acceptor scores in genes optimized to have high mRNA secondary structure. **J, K)** Correlations between gene lengths and the SpliceAI maximum donor or acceptor scores in genes optimized by Linear Design to have a high Codon Adaptation Index.

We then used our best performing model (RF_codons) to evaluate GFP sequences generated by various optimization tools (**Fig. 6B**). These optimized sequences were, on average, as similar to the training set as the test sequences used for model evaluation, indicating that predictions were made within the training distribution and are therefore likely reliable. Predicted expression varied widely across tools: JCat, GeneWiz/Azenta, LinearDesign and Twist produced sequences with expression levels comparable to the best variants in our library, while other designs showed poorer performance. Critically, nearly all tools—including the top performers—introduced cryptic splice sites with SpliceAI scores >0.2, suggesting a substantial risk of expression loss due to missplicing (Jaganathan et al., 2019). Some widely used platforms, including IDT, Optipyzer, and Basebuddy, generated sequences with splice site scores >0.7, indicating high susceptibility to splicing disruption.

Given that existing tools commonly generate sequences we predict to be misspliced, we evaluated several strategies for filtering away such sequences, using splicing data we collected for the 4,675 GFP variants as our benchmark. We sorted all variants based on their maximum SpliceAI or PSSM scores across the coding sequence - either for donor sites, acceptor sites, or the combination of both. We subsequently applied a conservative filtering approach by selecting the lowest-scoring 5% of variants for each metric, based on the premise that such filtering would reduce the likelihood of splicing. All six filtering strategies reduced the overall fraction of spliced transcripts (**Fig. 6C**). Notably, filtering by SpliceAI acceptor scores proved the most effective, with 65% of filtered variants showing no evidence of splicing and 98% exhibiting null to low splicing (<5%) (**Fig. 6D**). To assess whether this strategy compromises expression potential, we used the RF_codons model to estimate the expected protein output across the library. The SpliceAI-acceptor selection did not significantly reduce predicted expression levels, supporting its practical usability in design pipelines **(Fig. 6E)**.

To evaluate the risk of missplicing across a broader set of genes, we applied LinearDesign, an algorithm that performed well on our GFP benchmark, to 992 random human cDNAs from the MANE database. The resulting mRNA sequences were analysed using SpliceAI to predict the maximum splicing donor and acceptor probabilities across all positions of each gene. Most optimized genes had at least one position with a donor or acceptor score greater than 0.2, indicating a substantial risk of missplicing. Interestingly, donor and acceptor probabilities were lower in genes optimized to have high CAI than in the genes optimized to have high mRNA secondary structure **(Fig. 6 F, G).** Moreover, there was a negative correlation between gene lengths and SpliceAI scores, when genes were optimized to have either higher secondary structure or high CAI **(Fig. 6 H-K).** These results indicate that higher codon optimality, rather than stronger mRNA secondary structure, prevents the occurrence of missplicing. Unexpectedly, shorter genes are more susceptible to acquiring cryptic splice signals during codon optimization.

## DISCUSSION

Our experiments reveal a hierarchy of sequence-dependent control of heterologous gene expression in human cells, where GC content is the primary determinant of mRNA abundance, local sequence elements govern splicing status, and codon adaptation fine-tunes translation efficiency. These results align with the "unwanted transcript hypothesis," which proposes that mammalian codon usage coevolved with quality control systems that detect and suppress transcripts derived from viruses, transposable elements, or cryptic transcriptional events (Radrizzani et al.). Transcripts with low GC content, high CpG or UpA dinucleotide frequency, and a lack of introns are recognized as non-self and are targeted at multiple steps including transcription inhibition via HUSH (Seczynska et al.), nuclear degradation via NEXT and PAXT (Lubas et al. ; Giacometti et al.), nuclear export inhibition (Mordstein et al. ; Zuckerman et al.), sequestration in P-bodies (Courel et al.), and cytoplasmic decay mediated by ZAP and AU-binding proteins (Takata et al. ; Chen et al.). The GFP variants tested here, and many other intronless transgenes used in research and gene therapy, fall into the category of potentially "unwanted" transcripts and are therefore prone to these forms of surveillance. The redundancy of quality control pathways might explain why most codon effects are similar between cell types (**Fig. S20**), and why it is difficult to increase transgene expression through genetic manipulation of host cells.

The detection of unintended splicing in a substantial fraction of synonymous GFP variants highlights the sensitivity of transcript integrity to nucleotide-level changes. This observation aligns with previous findings that 20-60% of nucleotide changes within exons cause significant shifts in splicing patterns (Mueller et al. ; Julien et al. ; Savisaar and Hurst ; Baeza-Centurion et al. 2020). Clinically, a wide body of literature demonstrates that even single-nucleotide synonymous substitutions can significantly alter splicing in a plethora of genes, including *ATM*, *TP53*, *SMN2*, *APC*, and *NF1*(Sarkar, Panati, and Narala 2022). The *CFTR* gene exemplifies how single nucleotide changes can abolish or introduce splicing regulatory motifs, often with disease-relevant consequences (Pagani, Raponi, and Baralle 2005). Mechanistically, these disruptions stem from the creation or disruption of splicing regulators including exonic and intronic splicing enhancers and silencers (ESEs, ESSs, ISEs, ISS) that bind trans-acting factors with varying affinities, as well as canonical or cryptic splice sites. Ke et al. demonstrated through extensive mutagenesis that these elements function in a complex, interdependent network where context determines their activity (Ke et al. 2018). Beyond primary sequence, RNA secondary structure further modulates splicing by affecting the accessibility of binding motifs, as splicing regulators typically bind single-stranded regions, and stem-loop formations can sequester critical regulatory motifs (Buratti and Baralle 2004).

Despite the frequency and severity of these phenomena in natural genes, the literature includes only a few examples implicating recombinant DNA constructs (Dao et al. ; Paget-Bailly et al.). Epitope-tagged vectors such as those incorporating V5 tags have exhibited recurrent aberrant fusion transcripts due to activation of splice junctions at tag boundaries (Cheng et al. 2022; Ansseau et al. 2015). Additional reports reveal that even basic manipulations such as intron removal (wild type ORF expression) are sufficient to destabilize splicing both in human (Ono et al. 2022; Matsuoka et al. 2022) and non-human expression models (Top et al. 2021). Interestingly, a recent analysis of a codon-optimized SARS-CoV-2 spike gene revealed cryptic splicing that removed the transmembrane domain, generating a secreted variant potentially relevant to vaccine immunogenicity (Kowarz et al. 2022). This phenomenon may extend to other therapeutic genes, as similar cryptic splicing events have been documented in several commercially-developed recombinant constructs, highlighting the need for splicing-aware design approaches.

Given this growing evidence, predictive models such as SpliceAI have emerged as critical tools for identifying potential splicing disruptions (Canson et al. 2023). Massively parallel splicing assays demonstrated that SpliceAI outperforms other predictors in identifying splice-altering variants, with area-under-curve values of 0.98 (Smith and Kitzman 2023). Recent work by Wu et al. has established specific threshold values (≥0.3) to efficiently prioritize variants with high splicing risk (Wu et al. 2024). In our study, filtering by SpliceAI acceptor scores removed most heavily spliced constructs, demonstrating the feasibility of incorporating splicing-aware filters into the codon optimization workflow.

Our sequence-to-expression models account for 62–75% of the variance in GFP expression across the variant library. The models performed well when applied to highly divergent sequences and retained predictive power across cell types and expression modalities (mRNA versus protein). This robustness likely reflects the conserved nature of the sequence-dependent regulation in our experimental system. Training on global mechanistic features, rather than one-hot encoded sequence identities, may facilitate generalisation, consistent with (Shen, Kudla, and Oyarzun). However, the models are not expected to generalise to all contexts: codon-mediated regulation can be modulated by promoter identity (Zhou et al.), intron architecture (Mordstein et al.), transcriptional localisation (Palazzo and Kang), and RNA versus DNA delivery (Mordstein et al.), which might reweight or bypass certain regulatory features. Within standard mammalian expression systems, though, we expect the models to remain broadly applicable.

We therefore used our model to benchmark codon optimization algorithms. Despite similar goals, existing tools produced GFP variants with over 20-fold differences in predicted expression levels. Tools that maximized codon adaptation and GC content (e.g., JCat, GeneWiz/Azenta, Twist) generally yielded high-expression predictions, while those based on alternative principles—such as codon harmonization (CHARMING) or mRNA stability modeling (iCodon)—tended to underperform. The tissue-specific method CUSTOM also performed inconsistently: kidney-optimized variants achieved moderate expression, whereas lung-optimized versions performed poorly, even when evaluated using lung- and kidney-specific data from A549 and HEK293 cells, respectively. In addition, most models, including the top-performing ones, generated sequences predicted to cause some level of missplicing. Our benchmarking data is based on a single reporter gene, GFP, and more experiments are needed to assess generalisability. Nevertheless, our findings suggest that many optimization approaches overlook sequence features critical for transcript yield and fidelity in mammalian cells. Optimization strategies that explicitly account for RNA surveillance and splicing will be essential for improving the reliability of synthetic gene expression.

Collectively, these findings reveal that codon optimization—while valuable for expression—can compromise transcript integrity by altering splicing events that have been refined by natural selection in endogenous genes. Our results and the supporting literature emphasize the necessity of integrating splicing prediction tools into routine design of synthetic constructs, particularly for applications where transcript integrity is crucial for transgene efficacy and safety.

## MATERIALS AND METHODS

### Library designs

We analysed five coding-sequence libraries spanning GFP, mKate, and human genes: the GFP_DOE library (194 variants), GFP_optimized library (4,675 variants), mKate library (27 variants), ORFeome library (6,002 genes), and Therapeutic library (28 constructs). Together, these collections were used to dissect how synonymous sequence features influence expression, stability, and splicing across a range of sequence contexts.

The GFP_DOE library comprised 194 GFP variants designed to sample combinations of GC content, CpG frequency, and codon usage within a controlled amino acid sequence. A total of 168 variants were synthesized by GeneArt (Thermo Fisher Scientific) as described in (Notka, Liss, and Wagner) and cloned into Gateway entry vectors pENTR221 or pGK3 (Kudla et al.), and 31 variants from (Mordstein et al.) were cloned into pGK3. One variant carrying a nonsynonymous substitution and four variants that could not be cloned were excluded, yielding 194 constructs. This library was arrayed, with each variant cloned and sequence-verified individually, and was analysed either as individual constructs or in pooled format depending on the assay.

The GFP_optimized library was generated by introducing degenerate third-position substitutions across codons. For amino acids with six synonymous codons, only the fourfold-degenerate subset was used; together with other fourfold families, these positions were encoded as XXN with nucleotide frequencies of 40% G, 40% C, 10% A, and 10% T. For twofold families, two designs were used: XXR (R = 80% G, 20% A) and XXY (Y = 80% C, 20% T).

Isoleucine was encoded by ATY, and methionine and tryptophan by their single codons. This design yielded an average pairwise Hamming distance of 85 silent substitutions, an average GC3 of 84% and an average CAI of 0.75. Overlapping oligonucleotides were synthesized and assembled into full-length GFP genes by polymerase chain assembly, quantified by qPCR (170 fmol), and amplified (17 fmol input) before BamHI/EcoRI ligation into pENTR221 and E. coli transformation (∼10⁶ cfu). All colonies were pooled for plasmid preparation, and this library was maintained and analysed as a pooled collection. Following long-read sequencing, 4,675 constructs from this library passed the quality cutoffs of having no amino acid substitutions and at least 20 reads per variant.

The mKate library comprised 27 synonymous mKate variants designed in-house, spanning GC3 from 26% to 99%. Each variant was cloned and tested individually in an arrayed format.

The ORFeome library consisted of 6,002 intronless human open reading frames (ORFs) from 5,408 genes, obtained from the MGC Premier Human ORFeome Collaboration Collection (Stratech, UK). Each sequence included a terminal stop codon. This library was analysed as a pooled collection; splicing data were obtained for 689 of the 6,002 constructs with at least 20 reads per variant.

The Therapeutic library was provided by the U.S. Food and Drug Administration (CBER) and comprised 28 cDNA constructs representing clinically relevant proteins—ADAMTS13, MDTCS, and coagulation factors VIII and IX. For each gene, wild-type and optimized designs were included, representing codon-optimized, codon-pair-optimized, CpG-depleted, and other engineered variants. Codon optimization was performed using both commercial platforms (e.g., GenScript) and in-house algorithms targeting defined sequence features, including a codon-pair optimization method based on (Lin et al.). All constructs were cloned individually.

### Vector and barcoding systems

Two barcoded mammalian expression vectors were developed for library cloning and pooled analyses: pLCA1 and pNB1/BC, both derived from pcDNA5/FRT/TO/ccdB (a kind gift from Aleksandra Helwak). Each vector retains the FRT site for Flp-In–mediated genomic integration and the ccdB Gateway cassette for LR recombination with entry clones carrying individual inserts.

pLCA1 was generated by inserting a 20-nt random barcode into the 3′ untranslated region (UTR) of the pcDNA5/FRT/TO/ccdB expression cassette. Restriction sites NheI and KasI were first introduced by site-directed mutagenesis and sequence-verified in *E. coli* ccdB Survival 2 T1 cells (Thermo Fisher Scientific). The barcode insert, a 120-bp oligonucleotide containing a 20-nt degenerate region (IDT), was converted to double-stranded DNA by a single PCR cycle with Phusion polymerase, purified, and digested with AvrII and KasI. The vector backbone was digested with NheI-HF and KasI and dephosphorylated with Antarctic phosphatase. Barcoding was performed in large-scale ligations using 1 µg of vector and a 1:5 vector:insert molar ratio in a 3 mL total volume (10 µL T4 DNA ligase, 300 µL ligase buffer, 20 µL 100 mM ATP), incubated overnight at 16 °C. Pooled ligations were digested with NheI-HF for 1 h at 37 °C to remove recircularized vector and transformed into ultracompetent *E. coli* ccdB Survival 2 T1 cells prepared by the Inoue method. Approximately 1 × 10⁵ colonies were obtained, scraped into cold LB, pooled, and plasmid DNA was purified using the Qiagen Plasmid Maxi Kit. The resulting pLCA1 vector supports both transient transfection and site-specific integration into Flp-In host cells.

pNB1/BC was constructed as a modified pcDNA5/FRT/TO/ccdB backbone containing two additional tetracycline-responsive elements, a shortened 3′ UTR, and longer 30-nt barcodes. A multiple-cloning site was introduced at the PspOMI site using annealed oligonucleotides, and the TET repressive elements and 3′ UTR modifications were introduced sequentially by site-directed mutagenesis, with each step sequence-verified, yielding the pNB1 plasmid. For barcoding, the pNB1 plasmid was digested with NheI-HF and AscI and dephosphorylated with Shrimp Alkaline Phosphatase. The barcode duplex (30-nt degenerate insert) was generated from oligos NB0029 and NB0030 (1:2.5 ratio), annealed, redigested, and purified. Ligation used 85 ng total DNA at a 1:10 vector:insert ratio in 50 µL with T4 DNA ligase, incubated overnight at 4 °C. The product was treated with ClaI and KasI to reduce background, purified, and electroporated into *E. coli* ccdB Survival 2 T1 cells made electrocompetent by the Inoue method. Ten parallel 0.1 cm cuvettes (∼90 µL each) were pulsed at 2.5 kV, 200 Ω, 25 µF, and immediately recovered in pre-warmed SOC. Cultures were pooled, grown for 1 h at 37 °C (250 rpm), diluted to 1 L LB, and plated to estimate library complexity. Transformations yielded ∼1.26 × 10⁷ CFU, with ∼1% background from no-barcode controls, corresponding to ∼1.24 × 10⁷ unique barcoded plasmids. The pooled culture was expanded at 18 °C to OD₆₀₀ ≈ 1 and plasmid DNA was prepared using Qiagen miniprep columns to generate the pNB1/BC backbone for pooled high-diversity assays.

### Cloning and barcode–insert phasing

Coding sequences were transferred from Gateway entry vectors into barcoded expression vectors using LR Clonase II (Invitrogen). Each 10 µL reaction contained 100 ng of entry vector, 100 ng of destination vector, and 1 µL LR Clonase II, incubated for 20 h at room temperature. Reactions were transformed into E. coli DH5α, plated on LB–ampicillin, and incubated overnight at 37 °C. For the GFP_DOE library, individual variants were sequence-verified, transferred into the destination vector, purified, and combined in equimolar amounts to yield an expression-vector pool with equal representation of variants. For the other libraries, variants were transferred into the destination vector either as a single pool or as several smaller pools. The GFP_DOE and GFP_optimized libraries were cloned into pLCA1 (20-nt barcodes), while the mKate and ORFeome libraries were cloned into pNB1/BC (30-nt barcodes). The Therapeutic library was provided as individual expression plasmids and transfected directly, without additional barcoding.

Barcode–insert relationships were established by long- or short-read sequencing depending on the library. GFP_DOE phasing was determined using both Illumina (a 726-bp amplicon sequenced on MiSeq, 2 × 150 bp) and Oxford Nanopore (a KasI-digested 938-bp GFP–barcode fragment). The Nanopore library was prepared without PCR using the ligation sequencing kit (SQK-LSK110) and sequenced on a MinION instrument with R9.4.1 flow cells (FLO-MIN106). The Nanopore data were used to verify equal representation of variants across the library. The GFP_optimized library was phased using PacBio long-read sequencing, and the mKate and ORFeome libraries were phased by Oxford Nanopore sequencing and read mapping with minimap2. In all cases, barcode sequences were clustered (≤ 2 mismatches) and retained if supported by at least two reads and uniquely associated with a single variant. The Therapeutic constructs lacked barcodes; instead, multiplexed RT–PCR with primer-specific barcodes allowed direct assignment of cDNA reads to the corresponding plasmid.

### Cell lines and transfection protocols

HeLa and HEK293 cells were obtained from ATCC. HEK293 ZAP⁻/⁻ and isogenic wild-type cells were described previously (Sharp et al.). HeLa TRIM25⁻/⁻ cells and their parental controls were a gift from Gracjan Michlewski and Nila Chowdhury (Choudhury et al.). SHSY5Y, A549, HepG2, and hTERT RPE1 cells were provided by Cathy Abbott, Lesley Stark, Nick Gilbert, and Duncan Sproul (Institute of Genetics and Cancer). All cell lines were maintained at 37 °C and 5% CO₂.

Growth media and exact transfection formats, DNA amounts, and reagent ratios for pooled and arrayed experiments are listed in Table S1. Unless stated otherwise, cells were seeded one day prior to reach ∼60–70% confluency and harvested 24 h after transfection.

### RNA quantification, mRNA half-life, and serum-withdrawal assays

To quantify steady-state mRNA levels, the pooled GFP_DOE library was transfected into HEK293, HeLa, A549, SHSY5Y, HepG2, and hTERT RPE1 as per Table S1. Twenty-four hours post-transfection, total RNA was extracted (RNeasy, Qiagen), treated with RNase-free DNase, and rRNA was depleted (NEB). Libraries were prepared without PCR using the Oxford Nanopore ligation kit v14 (SQK-LSK114) and sequenced on R10.4.1 flow cells (MinION/PromethION), preserving relative transcript abundances.

For mRNA half-life measurements, HeLa cells were transfected with the pooled GFP_DOE library; transcription was inhibited 24 h later with 500 nM Triptolide, and RNA was collected at 0, 1, 2, 4, 6, and 8 h. RNA was processed as above and sequenced using the same PCR-free ligation workflow.

For serum withdrawal, HeLa cells were transfected with the pooled GFP_DOE library, switched to serum-depleted medium for 24 h, and RNA was isolated and sequenced using the same PCR-free ligation protocol.

As an orthogonal control for RNA abundance, arrayed qRT-PCR assays were performed on individual GFP_DOE constructs: HeLa cells were reverse-transfected (Table S1), RNA was isolated (RNeasy or Trizol), DNase-treated, and reverse-transcribed with SuperScript III; qPCR used SYBR Green I with primers against GFP and NeoR, and relative expression was calculated by 2^–ΔΔCt with NeoR normalization. Data processing and quantification for all RNA assays are described under Computational analysis.

### Splicing assays

Cryptic splicing was quantified in HeLa after pooled transfection of the GFP_DOE, GFP_optimized, mKate, and ORFeome libraries; Therapeutic constructs were transfected individually and pooled when collecting cells. For pooled libraries only, cycloheximide was added at 20 h. Twenty-four hours post-transfection, total RNA was extracted, DNase-treated, and rRNA-depleted. RNA was reverse-transcribed; a 9-nt UMI was appended in a single-cycle indexing PCR, followed by 18 PCR cycles for amplification. Libraries were sequenced on Oxford Nanopore instruments (MinION Mk1C) as appropriate for each dataset. GFP_DOE phasing was additionally verified by Nanopore sequencing of KasI-digested plasmid amplicons (938 bp) prepared without PCR using the ligation kit SQK-LSK110 and MinION R9.4.1 flow cells (FLO-MIN106). Data processing, barcode assignment, and isoform quantification are described under Computational analysis.

### Protein expression assays

For transient protein measurements, individual GFP_DOE variants were transfected in triplicate into HeLa, HEK293, and A549 in 96-well plates (Table S1). Before the transfection, 96-well plates were coated with Poly-L-Lysine to promote the adhesion between cells and the plate surface. Twenty-four hours post-transfection, media was removed and cells were lysed by adding 200 µL of cell lysis buffer (25 mM Tris, pH 7.4, 150 mM NaCl, 1 % Triton X-100, 1 mM EDTA, pH 8) to each well. Cell lysis was performed before fluorescence measurements to prevent fluctuations in fluorescence intensity due to uneven distribution of cells within the wells. GFP fluorescence was measured on a Tecan Infinite M200 Pro (Ex 486 nm / Em 515 nm). Background from non-transfected wells was subtracted, and across-plate normalization used three shared GFP controls (low/medium/high GC).

ZAP/TRIM25 dependency was assessed by transfecting a subset of GFP variants into isogenic wild-type and knockout (ZAP⁻/⁻ or TRIM25⁻/⁻) lines using the same arrayed protocol. Twenty-four hours later, lysates were assayed for GFP and mKate2 fluorescence (Ex 486/Em 515; Ex 588/Em 633), and GFP was normalized to mKate2 per well. Statistical analyses are detailed under Computational analysis.

For stable expression profiling (Flow-seq), pooled barcoded GFP_DOE constructs were integrated into HeLa Flp-In T-REx cells by co-transfection of pcDNA5/FRT/TO-GFP-BC and pOG44 (1:9). Stable integrants were selected with blasticidin S (10 ng/μL) and hygromycin B (400 mg/mL) for 2–3 weeks until negative controls were cleared. Doxycycline (1 μg/mL) induced expression for 24 h before sorting on a BD FACS Aria II into eight fluorescence bins. Genomic DNA from each bin was used to prepare Illumina amplicon libraries (NextSeq PE75). Barcode mapping and fluorescence-coefficient calculations are described under Computational analysis.

### Computational analysis: data processing and quantification

Barcode–insert phasing was established prior to downstream analyses. GFP_DOE barcodes were linked to coding sequences using both Illumina (MiSeq 2 × 150 bp) and Nanopore (MinION R9.4.1, SQK-LSK110) sequencing of plasmid libraries. GFP_optimized constructs were phased using PacBio long-read sequencing, while mKate and ORFeome libraries were phased with Oxford Nanopore (SQK-LSK114). Illumina and Nanopore reads were basecalled, filtered, and mapped to plasmid references using minimap2 (v2.16). Mapped reads were converted to FASTQ, trimmed to extract barcode regions with cutadapt (v1.9.1), and clustered using cd-hit-est (≤ 2 mismatches). Only clusters supported by at least two reads and uniquely associated with a single variant were retained to generate verified barcode–variant reference tables.

Nanopore and Illumina sequencing data from RNA expression, half-life, serum-withdrawal, and splicing assays were processed through a unified pipeline. Raw FAST5 files were basecalled with Guppy (v6.5.7) and filtered with Nanofilt to retain reads with Phred quality scores above 10. Filtered FASTQ reads were aligned to either the transgene+barcode reference (for GFP and mKate datasets) or the MANE human transcriptome (for ORFeome and Therapeutic libraries) using minimap2. Primary alignments (SAM flags 0 and 16) with MAPQ > 10 were extracted with samtools (v1.10) and visualised in IGV. Barcode-level counts were obtained by summing reads assigned to each construct. Equivalent mapping parameters were used for Nanopore DNA sequencing of plasmid libraries.

Splicing assays were processed from PCR-amplified cDNA libraries containing 9-nt UMIs. Reads were demultiplexed by barcode, and unique isoforms were identified from CIGAR strings in minimap2 alignments: the number of ‘N’ operations defined the intron count, intron lengths were taken from the numeric values preceding each ‘N’, and exon block lengths were computed from contiguous M/D stretches. Only barcodes or variants with at least 20 UMI-unique reads were retained. Splicing fraction per variant was calculated as the proportion of UMI-unique reads containing at least one intron.

Flow-seq libraries were quantified from Illumina NextSeq PE75 datasets. Barcode counts per fluorescence bin were used to calculate a fluorescence coefficient (FC) as the weighted mean of bin fluorescence intensities, using the median fluorescence per bin and read count as weights.

For mRNA half-life assays, transcript abundances over time were fitted to an exponential decay model R(t) = R₀·e^–λt. Fits and half-lives (t₁/₂ = ln2/λ) were obtained by non-linear regression in Python. Only transcripts with ≥10 reads at time 0, half-lives between 0–20 h, and R² ≥ 0.2 were retained. Translation efficiency for each construct was calculated as the ratio of its protein to mRNA expression.

### Computational analysis: sequence feature computation

GC3 content was calculated from coding sequences using custom awk scripts that parsed codons, extracted the third nucleotide, and computed the fraction of G or C. CpG counts were determined as the total number of CpG dinucleotides within each coding sequence. Codon adaptation index (CAI) was calculated with EMBOSS (v6.6.0) using codon usage from highly expressed human genes as defined by (Mordstein et al.). Codon frequencies for each variant were correlated with gene expression to derive codon expression coefficients (CECs) using Pearson’s correlation in R.

In-silico scoring of splicing-related features was performed on full amplicon sequences extending from the 5′UTR (233 nt upstream of the start codon) through the CDS and 94 nt into the 3′UTR, just upstream of the barcode. Donor and acceptor position-specific scoring matrices (PSSMs) were constructed from ∼90,000 annotated human splice junctions, using windows of [–3, +6] around GT donors and [–14, +1] around AG acceptors, and branchpoint scores were taken from the BPP repository. For each acceptor site, the maximum branchpoint score within – 80 to –15 nt and the maximum polypyrimidine tract (PPT) score within –45 to –1 nt were recorded. PPT strength was quantified using sliding windows of 15 or 20 nt by computing (i) the proportion of pyrimidines (C+T), (ii) the proportion of T alone, and (iii) a weighted pyrimidine ratio (T = 0.666, C = 0.333), with the maximum score across the window retained as the PPT value.

G-quadruplex propensity was estimated using G4Hunter (25-nt window, default parameters), recording the maximum value within ±100 nt of each splice site. Exonic splicing enhancer motifs were identified using the RESCUE-ESE database of 238 hexamers (Fairbrother et al.). SpliceAI (v1.3) was run in sequence mode, and donor and acceptor probabilities were extracted for all bases. Per-position scores were stratified by observed splice usage (0, >0–0.1, 0.1–0.9, >0.9), and per-construct analyses retained the maximum value across the full amplicon.

### Computational analysis: modeling and statistics

Predictive models were implemented in Python using scikit-learn. Three regression approaches were evaluated: linear regression (ordinary least squares), quadratic polynomial regression (degree-2 feature expansion followed by least squares), and random forest regression (200 estimators, maximum depth 8, minimum samples per split 5, minimum samples per leaf 4, log₂ feature selection, bootstrap disabled). Ten feature sets were defined, comprising either single global features (GC3, CpG, CAI, mRNA folding energy), codon-level features, splicing-related scores, or their combinations. Each model–feature set combination was trained to predict a specified expression variable, yielding 30 distinct runs.

Model performance was estimated using five-fold cross-validation (RepeatedKFold, 5 splits, 1 repeat, random state 42). For each fold, 80% of data were used for training and 20% for validation. Predictions on held-out subsets were combined to form an out-of-fold (OOF) prediction vector per construct. We computed (i) the mean and standard deviation of fold-wise R² values, (ii) pooled OOF R² across the dataset, and (iii) Pearson correlations (and squared correlations) between OOF predictions and measured values. The first two metrics describe within-modality performance, while squared correlations were used to compare cross-modality predictive power (e.g., protein versus RNA).

For benchmarking, a low-expression GFP variant (GFP_407) was re-optimized and de-optimized using publicly available gene design tools with default parameters. Expression predictions were obtained using a random forest model trained on the A549 GFP protein dataset (80/20 train–test split).

To test general sequence optimization strategies, 992 coding sequences from the Matched Annotation from NCBI and EMBL-EBI (MANE) transcript database were optimized with LinearDesign to maximize either mRNA secondary structure stability or codon adaptation. Genes were filtered to retain those with canonical start and stop codons, translated with EMBOSS transeq, and optimized with λ = 10 for high-codon adaptation runs.

Statistical dependencies between features were evaluated by partial correlation analysis in R. Expression values were first regressed against GC3 to obtain residuals; CpG or CAI values were likewise regressed against GC3, and Pearson correlations were computed between the residuals to estimate associations independent of GC3 effects.

## ACKNOWLEDGMENTS

GK was supported by Wellcome Trust (fellowship 207507) and the Medical Research Council (MRC) Human Genetics Unit core grant MC_UU_00035/8. MA was supported by the "Context-Dependent Gene Optimization" grant from Thermo Fisher Scientific GENEART GmbH. EG was supported by a Wellcome Trust Sir Henry Dale fellowship (211222/Z/18/Z) and a BBSRC Roslin Institute Strategic Programme Grant (BB/X010937/1). SW was funded by Lundbeck Foundation, the Novo Nordisk Foundation, Danish Cancer Association, Carlsberg Foundation, Independent Research Fund Denmark, Aage and Johanne Louis-Hansen’s Foundation.

## Declaration of generative AI and AI-assisted technologies in the writing process

During the preparation of this work the authors used the AI tool ChatGPT in order to improve language and readibility. After using this tool/service, the authors reviewed and edited the content as needed and take full responsibility for the content of the publication.

## FIGURE LEGENDS

**Supplementary figure S1.**
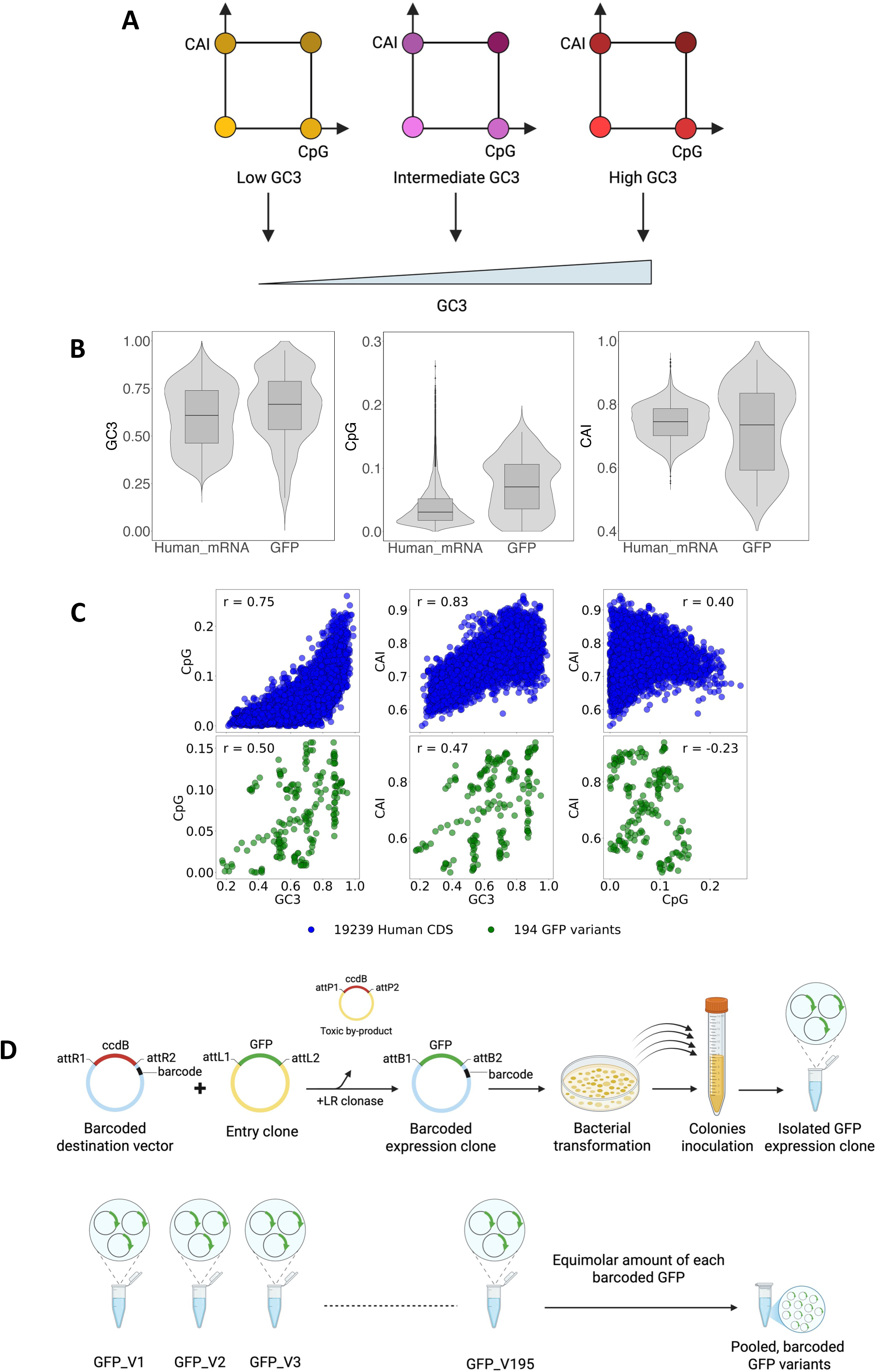
Design and evaluation of systematic codon analysis platform. **A)** Design of 96 GFP variants from the GFP_DOE library with constrained GC3 levels. **B)** Violin plots showing GC3, CpG, and CAI levels of human endogenous mRNAs and of the GFP library. **C)** Scatter plots showing correlation among the GC3, CpG, and CAI levels of endogenous human mRNAs (blue, n=19239) and GFP library (green, n=194). **D)** Schematic representation of the arrayed cloning strategy of the GFP_DOE library into a barcoded expression vector, followed by pooling of the library.

**Supplementary figure S2.**
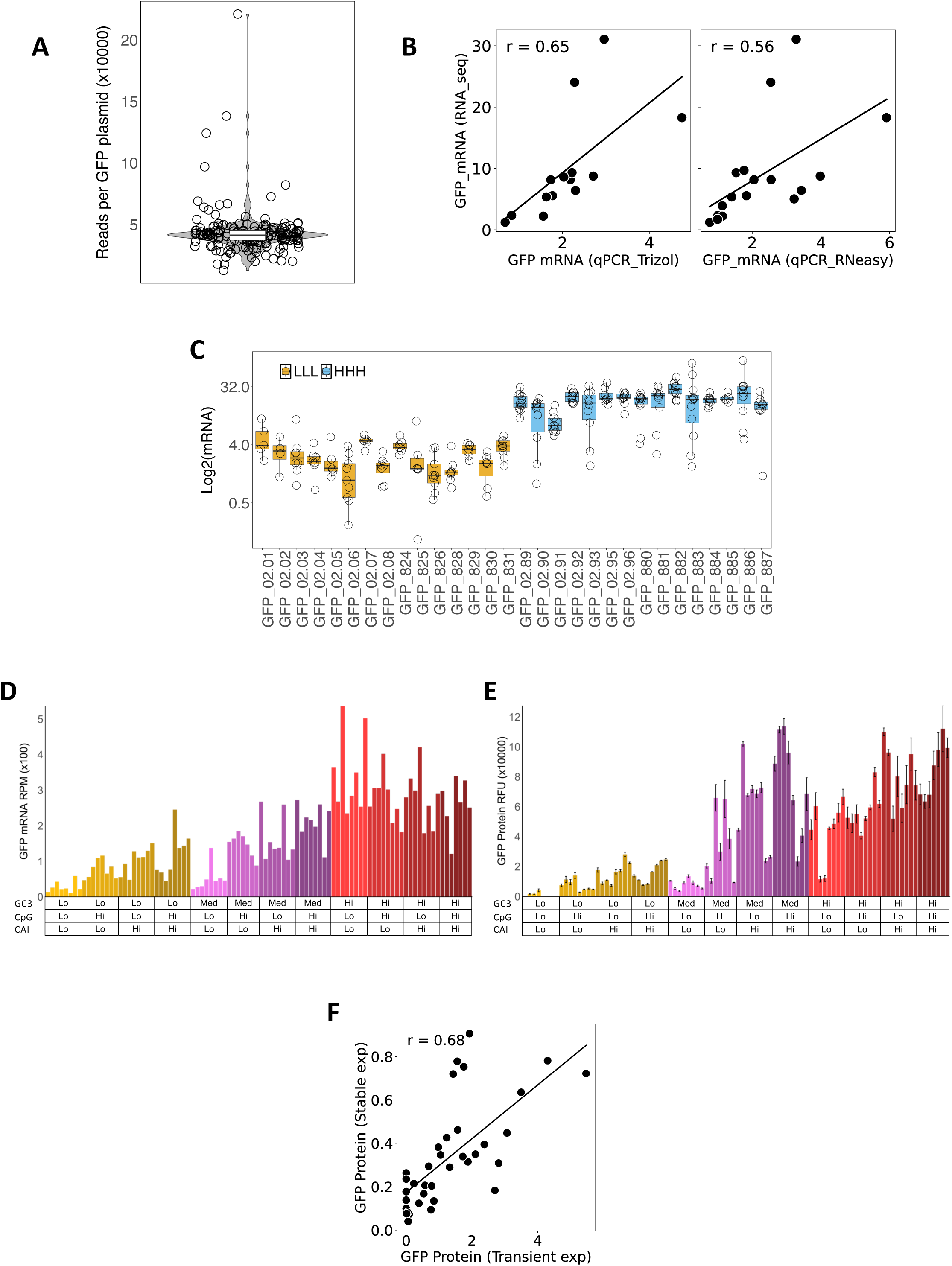
Preparation and quality control of GFP variant library. **A)** Violin plot showing the abundance of variants in the GFP_DOE library. **B)** Scatter plots showing correlations between GFP mRNA expression, measured by RNA seq, and the qPCR (using either Trizol or RNeasy method for total RNA isolation) in HeLa cells. **C)** Box plot showing the mRNA expression levels of the GFP variants in HeLa cells, according to the gene feature levels (LLL: low GC, low CpG, low CAI & HHH: high GC, high CpG, high CAI. Each circle is a barcoded replicate for the given GFP. **D, E)** Transient mRNA and protein expression of 96 variants from the GFP_DOE library in HeLa cells according to their gene feature levels in Figure S1A. **F)** Correlation between stable and transient GFP protein expression in HeLa cells. The x-axis shows GFP fluorescence intensity (×10⁴ RFU), and the y-axis shows GFP protein abundance measured by FlowSeq (×10⁴ AU).

**Supplementary figure S3.**
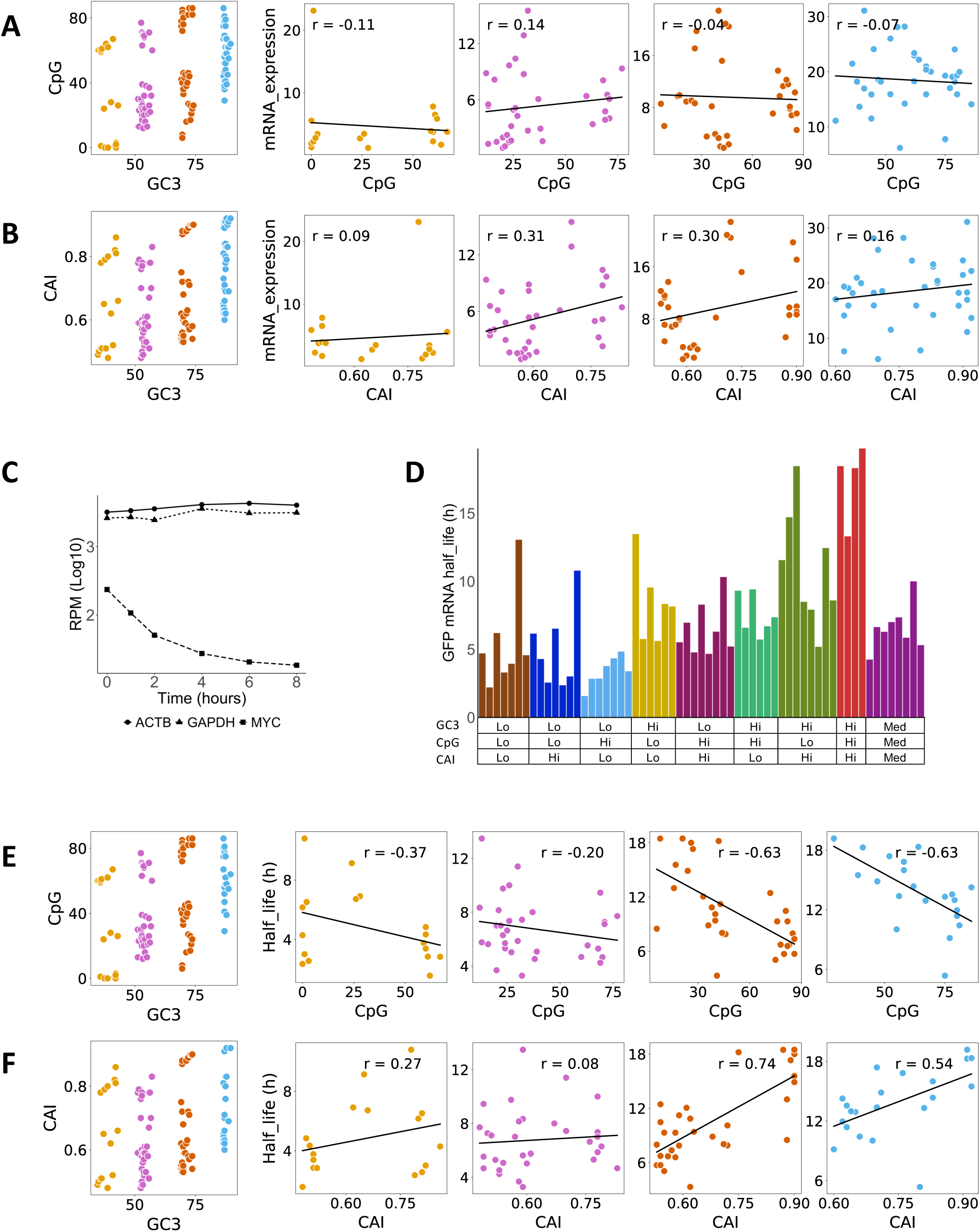
Effects of sequence properties on mRNA expression and mRNA half-life. **A, B)** GFP variants were stratified by GC3 content into four groups and each group’s CpG and CAI levels were plotted against mRNA expression in HeLa cells. **C)** Decay curve of stable mRNAs Actin beta and GAPDH, (half-life of >8h) and unstable mRNA c-myc (half-life of 0.96h). **D)** GFP mRNA half-lives of the 72 variants (Lo: Low, Med: Medium, Hi: High). **E, F)** GFP variants were stratified by GC3 content into four groups and each group’s CpG and CAI levels were plotted against the mRNA half-life in HeLa cells.

**Supplementary Figure S4.**
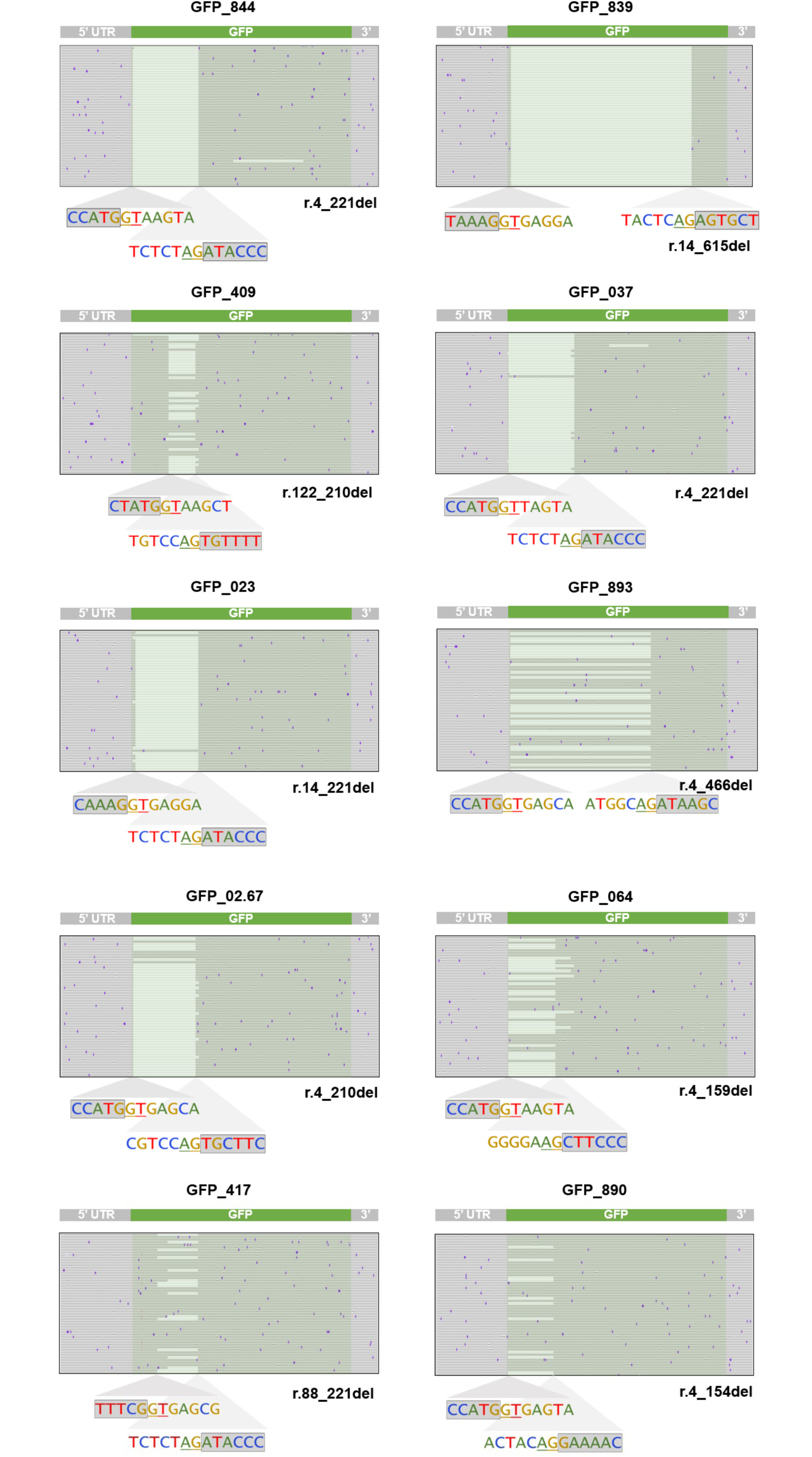
Splicing patterns of selected GFP variants in HeLa cells. IGV images showing missplicing and corresponding donor and acceptor sites across selected variants from the GFP_DOE library.

**Supplementary Figure S5.**
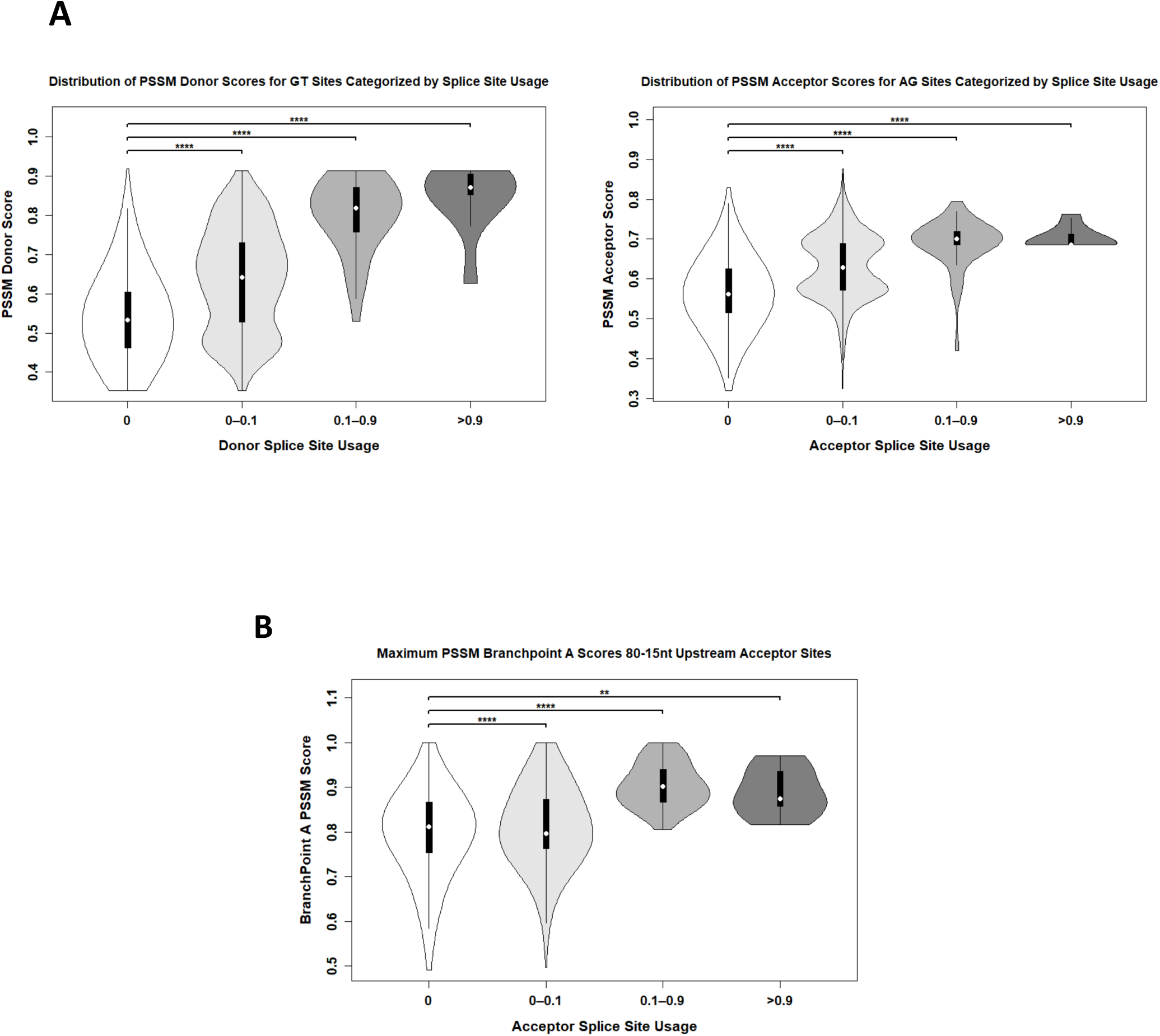
Correlation of PSSM acceptor scores, donor scores and branchpoint scores with splice site usage in the GFP_DOE library. **A)** Violin plots of donor and acceptor PSSM scores across all GFP variants at GT and AG positions in the GFP_DOE library, stratified by splice site usage. Significance was tested with Wilcoxon rank-sum tests (****p < 0.0001). **B)** Violin plots show the maximum PSSM Branchpoint A score within the −80 to −15 nt window upstream of each position, stratified by the observed splice acceptor usage of that position. Statistical comparisons were performed using Wilcoxon rank-sum tests. Significance: (p < 0.01 (**), p < 0.0001 (****), ns, not significant.

**Supplementary Figure S6.**
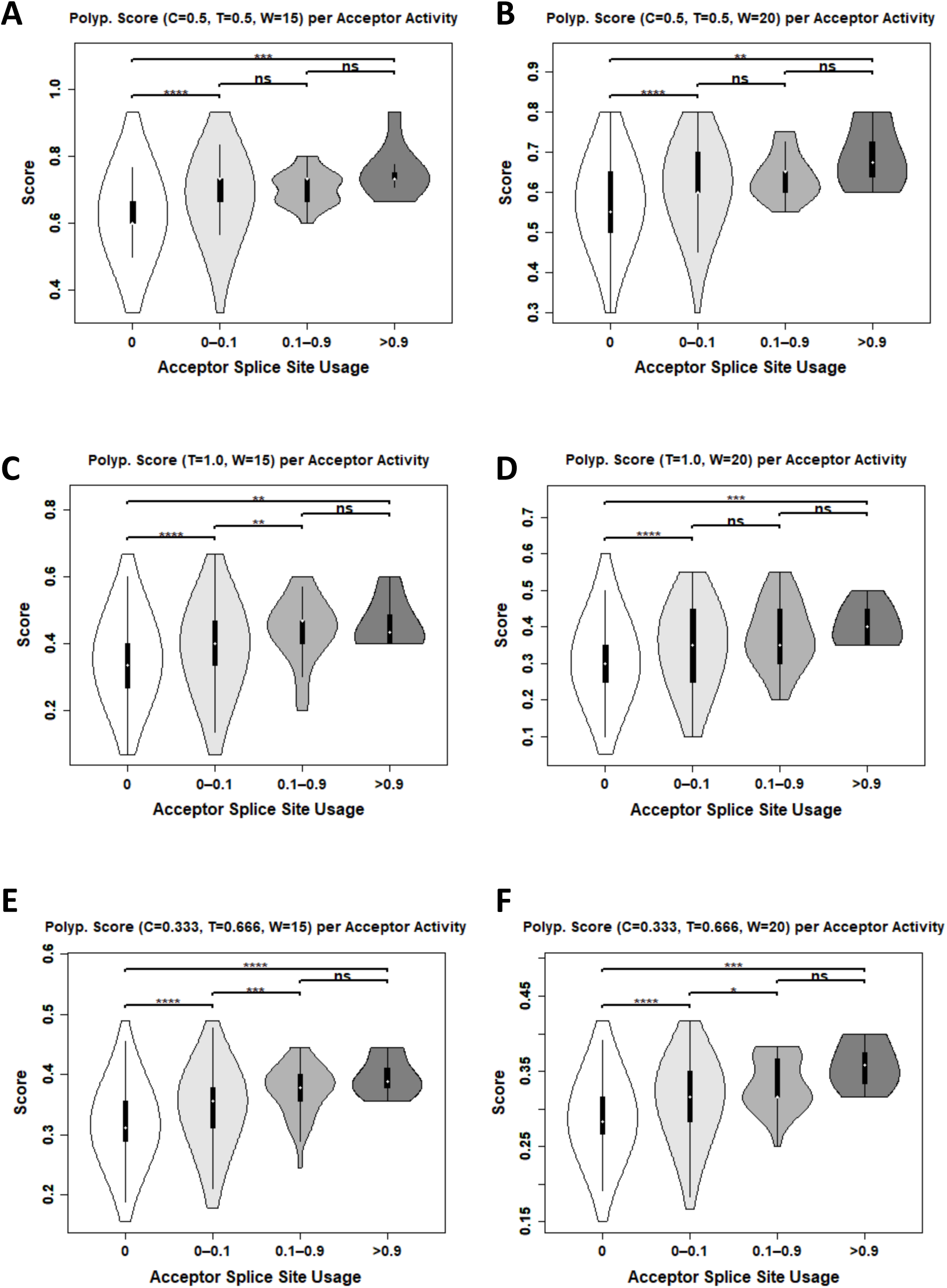
Correlation of polypyrimidine tract scores with splice acceptor site usage in the GFP_DOE library. (**A-F**) Violin plots show six different scoring strategies to evaluate polypyrimidine tract strength upstream of potential acceptor sites. For each nucleotide, the maximum score within a 45-nt upstream window was computed using: cytosine (C) and thymine (T) content with equal weight (C = 0.5, T = 0.5) in 15-nt **(A)** or 20-nt **(B)** windows; thymine-only scoring (T = 1.0) in 15-nt **(C)** or 20-nt **(D)** windows; and weighted C and T content (C = 0.333, T = 0.666) in 15-nt **(E)** or 20-nt **(F)** windows. Scores are stratified by acceptor splice site usage, revealing associations between polypyrimidine content and increased acceptor activity. Statistical comparisons were performed using a Wilcoxon rank-sum test. Significance: p < 0.05 (*), p < 0.01 (**), p < 0.001 (***), p < 0.0001 (****), ns, not significant. Polyp., polypyrimidine.

**Supplementary Figure S7.**
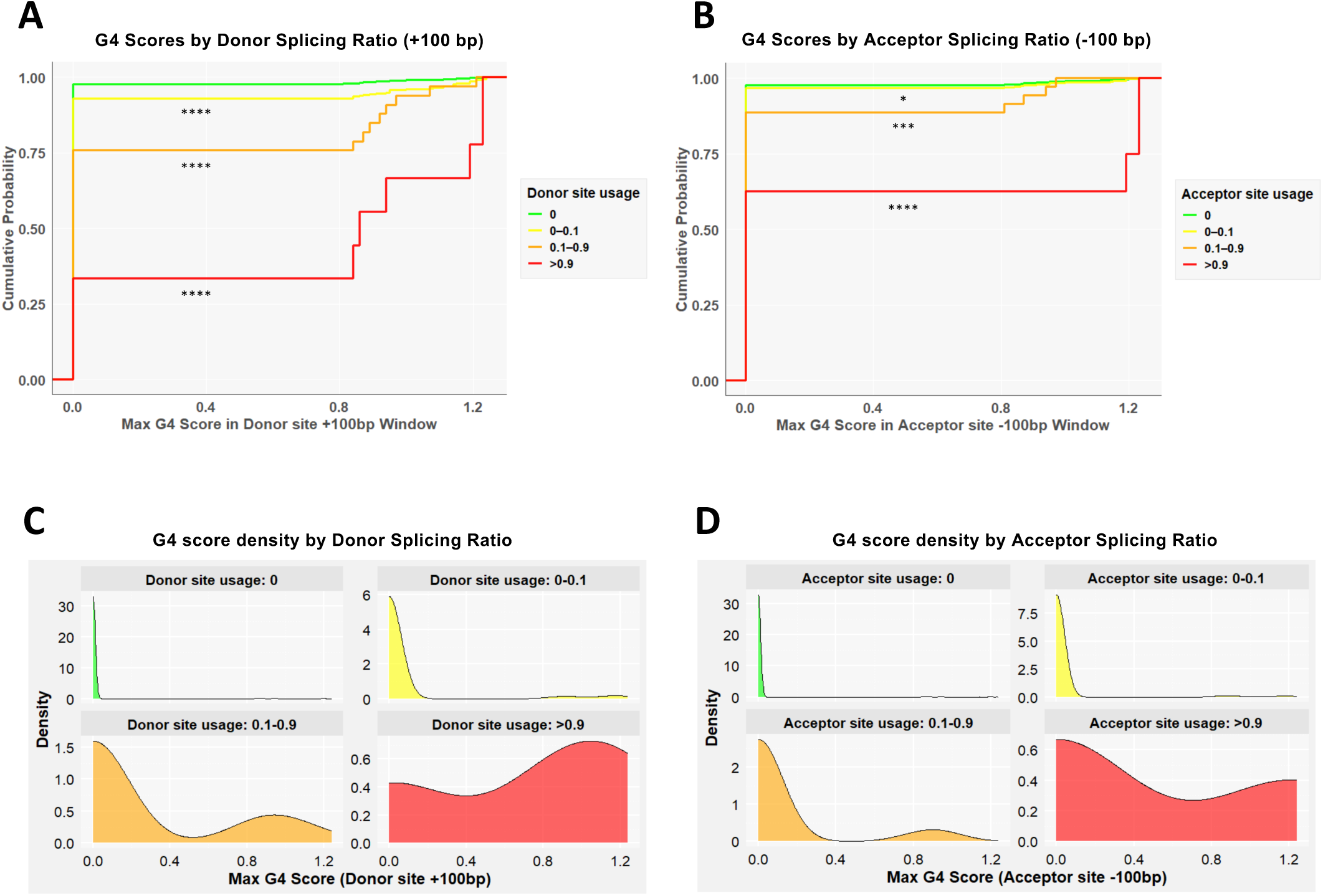
Correlation of G-quadruplex (G4) scores with splice site usage in the GFP_DOE library. To assess the influence of G-quadruplexes (G4) on splicing, we calculated G4Hunter scores at every nucleotide and recorded the maximum score within +100 nt downstream of each donor site or −100 nt upstream of each acceptor site. (**A, B**) Cumulative distribution plots show the maximum G4 scores stratified by splice site usage for donor (**A**) and acceptor (**B**) sites. Sites with high splicing activity (>0.9, red) show notable higher G4 scores for both donor and acceptor sites, revealing a modulatory role of G4 structures in splicing regulation. Significance: p < 0.05 (*), p < 0.001 (***), p < 0.0001 (****). (**C, D**) Density plots of the same G4 scores for donor (**C**) and acceptor (**D**) sites reveal distinct score distributions across splicing ratio groups. Curves shift rightward with increased splicing, suggesting higher G4 propensity at more frequently used splice sites.

**Supplementary Figure S8.**
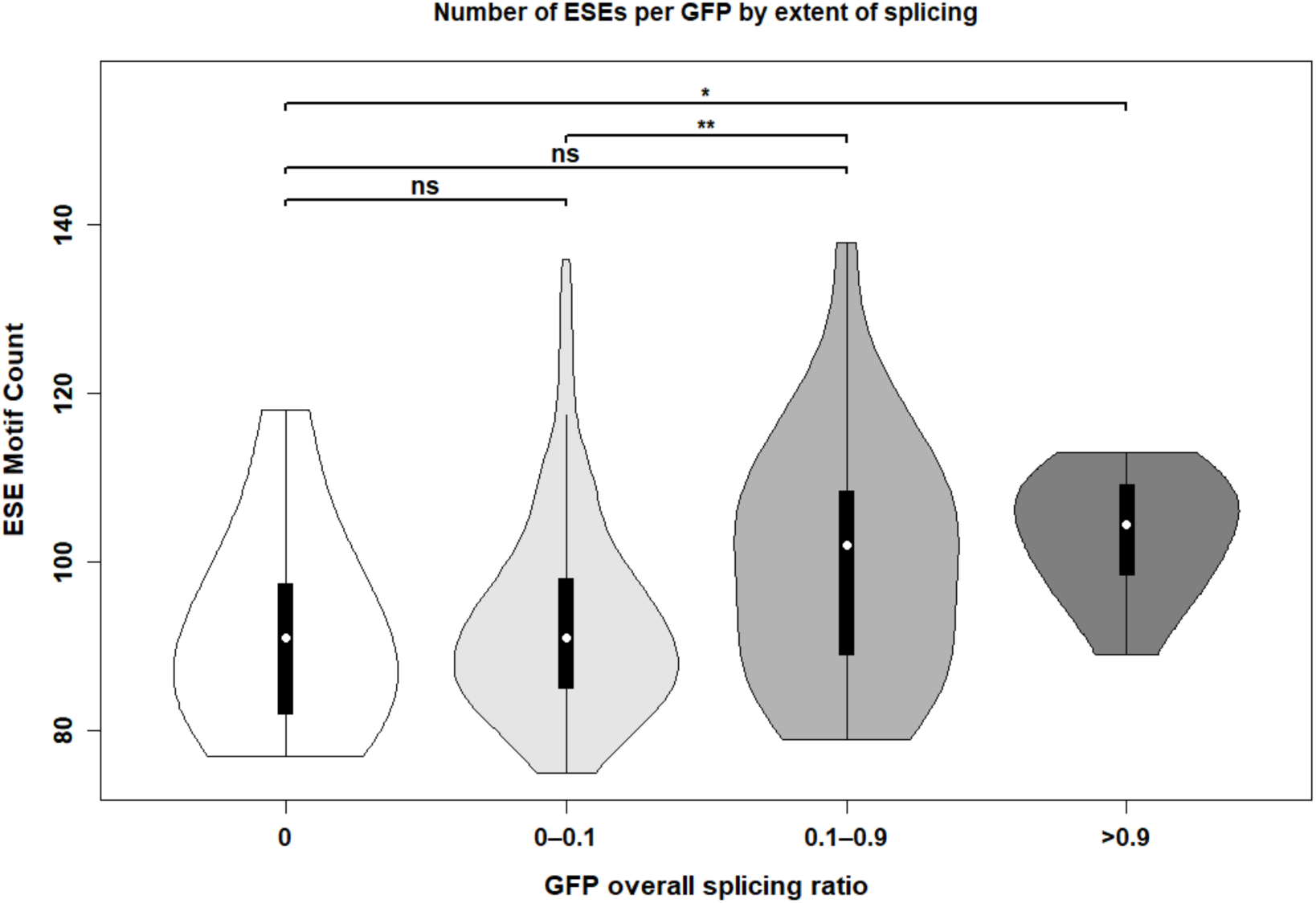
Exonic splicing enhancer (ESE) counts increase with overall splicing levels in the GFP_DOE library. Violin plots show the distribution of ESE counts across GFP variants grouped by their overall splicing ratio. The x-axis reflects bins of total splicing observed for GFP transcripts, while the y-axis indicates the number of identified ESE motifs per construct. A significant increase in ESE content is observed in highly spliced GFPs, showing that ESE abundance contributes to enhanced splice site recognition. Significance: p < 0.05 (*), p < 0.01 (**), ns, not significant.

**Supplementary Figure S9.**
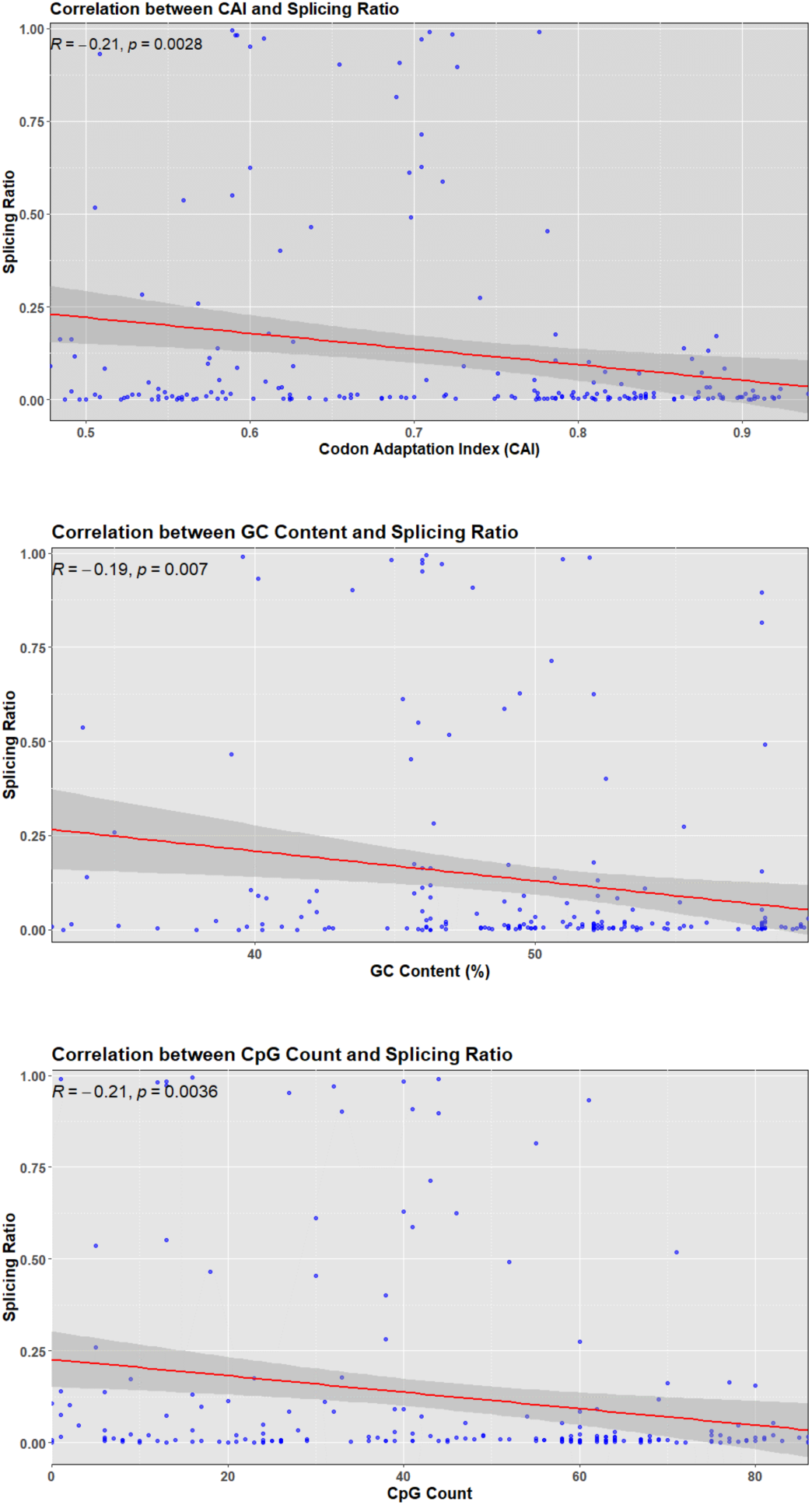
Correlation of Codon Adaptation Index (CAI), GC content and CpG count with splicing in the GFP_DOE library. Scatter plots show the relationship between the GFP mRNA splicing ratio and three sequence-derived features across all GFP variants: Codon Adaptation Index (CAI, top), GC content (middle), and CpG count (bottom). Each point corresponds to a single GFP variant. Linear regression (red line with 95% confidence interval in grey) reveals a weak but significant negative correlation between splicing and all three features: CAI (R = −0.21), GC content (R = −0.19), and CpG count (R = −0.21).

**Supplementary Figure S10.**
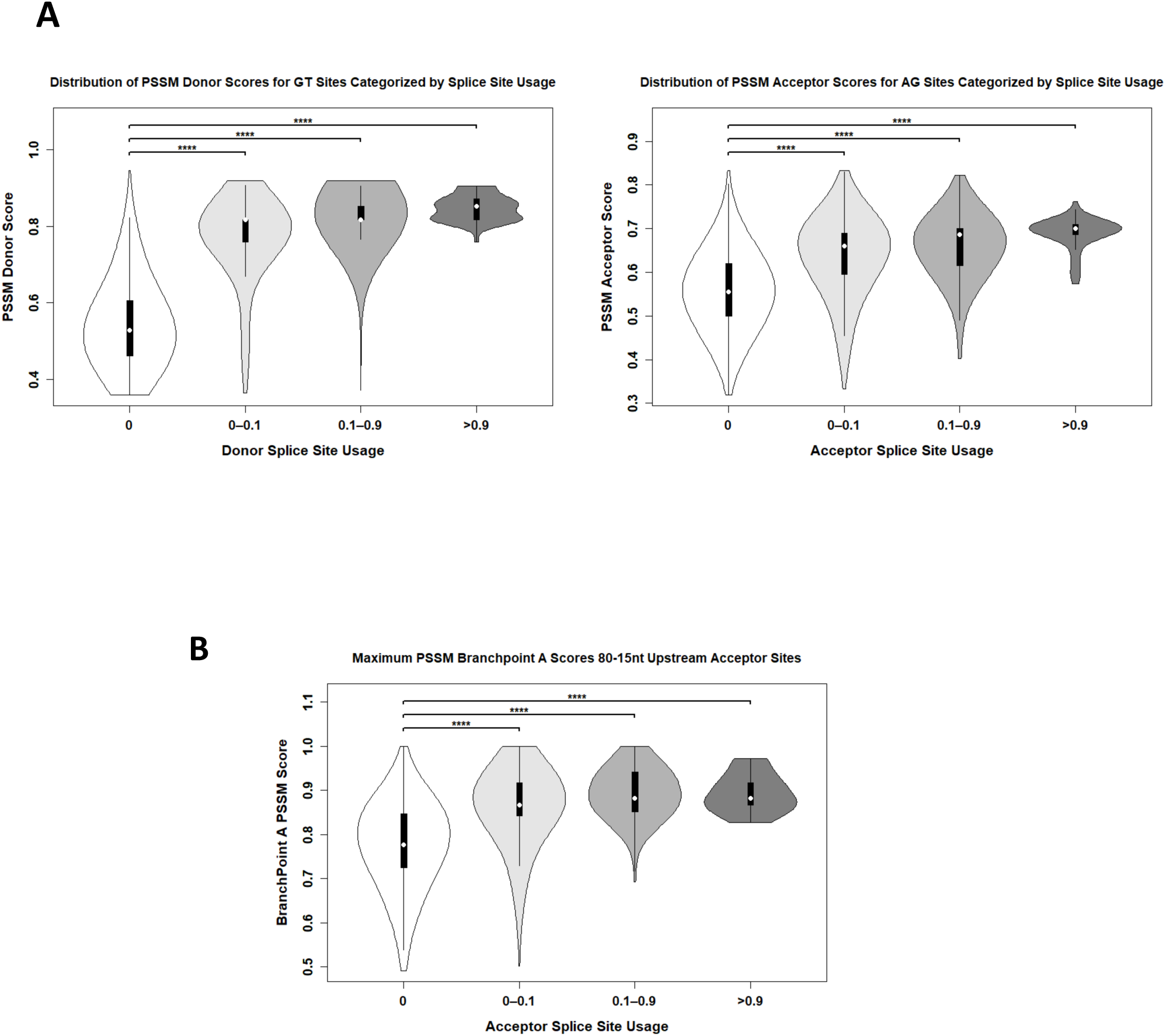
Correlation of PSSM acceptor scores, donor scores and branchpoint scores with splice site usage in the GFP_optimized library. **A)** Violin plots of donor and acceptor PSSM scores across all GFP variants at GT and AG positions in the GFP_DOE library, stratified by splice site usage. Significance was tested with Wilcoxon rank-sum tests (****p < 0.0001). **B)** Violin plots show the maximum PSSM Branchpoint A score within the −80 to −15 nt window upstream of each position, stratified by the observed splice acceptor usage of that position. Statistical comparisons were performed using Wilcoxon rank-sum tests. Significance: p < 0.0001 (****), ns, not significant.

**Supplementary Figure S11.**
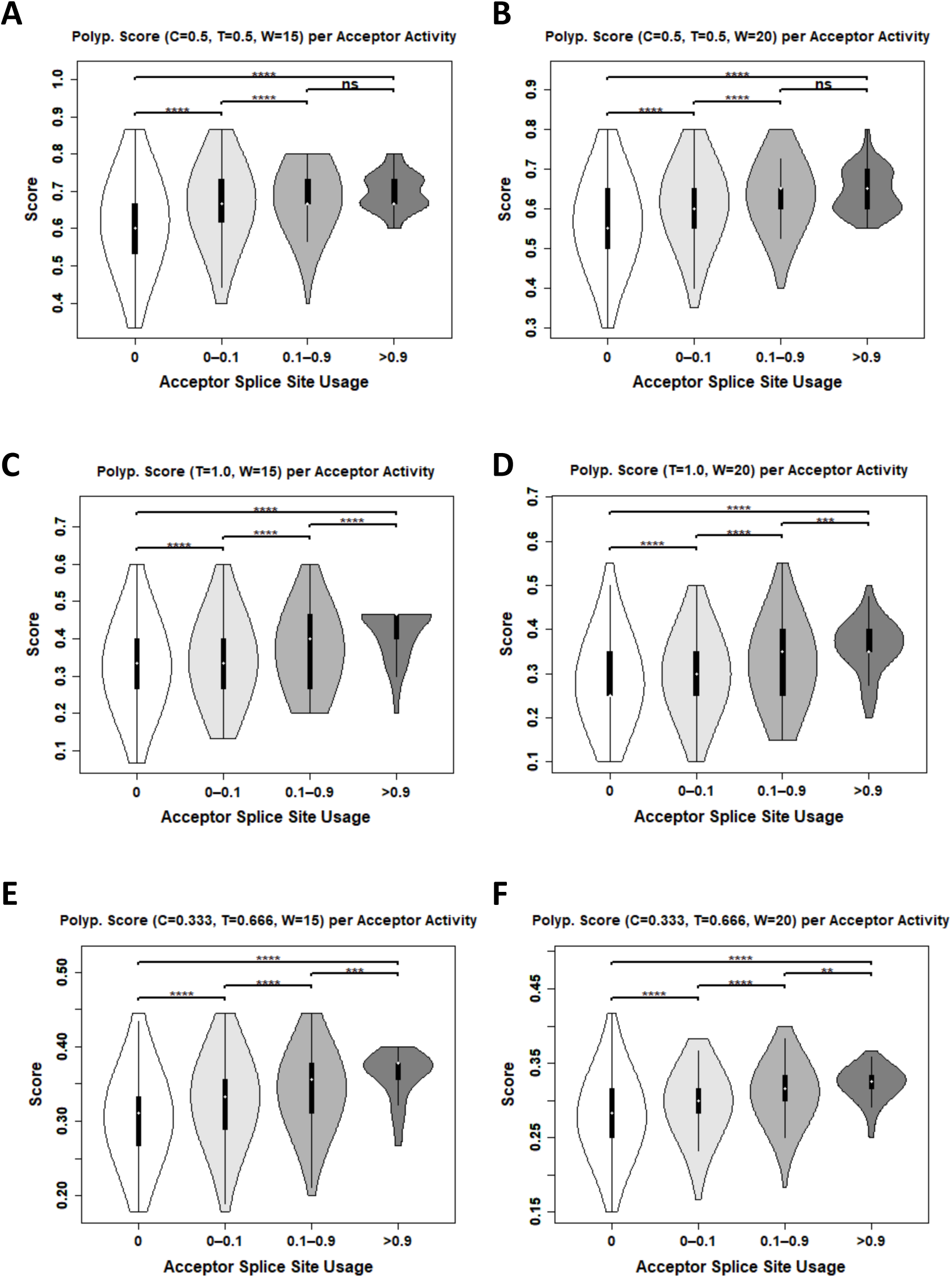
Correlation of polypyrimidine tract scores with splice acceptor site usage in the GFP_optimized library. Violin plots show six different scoring strategies to evaluate polypyrimidine tract strength upstream of potential acceptor sites. For each nucleotide, the maximum score within a 45-nt upstream window was computed using: cytosine (C) and thymine (T) content with equal weight (C = 0.5, T = 0.5) in 15-nt **(A)** or 20-nt **(B)** windows; thymine-only scoring (T = 1.0) in 15-nt **(C)** or 20-nt **(D)** windows; and weighted C and T content (C = 0.333, T = 0.666) in 15-nt **(E)** or 20-nt **(F)** windows. Scores are stratified by acceptor splice site usage, revealing strong and consistent associations between polypyrimidine content and increased acceptor activity. Statistical comparisons were performed using a Wilcoxon rank-sum test. Significance: p < 0.01 (**), p < 0.001 (***), p < 0.0001 (****), ns, not significant. Polyp., polypyrimidine.

**Supplementary Figure S12.**
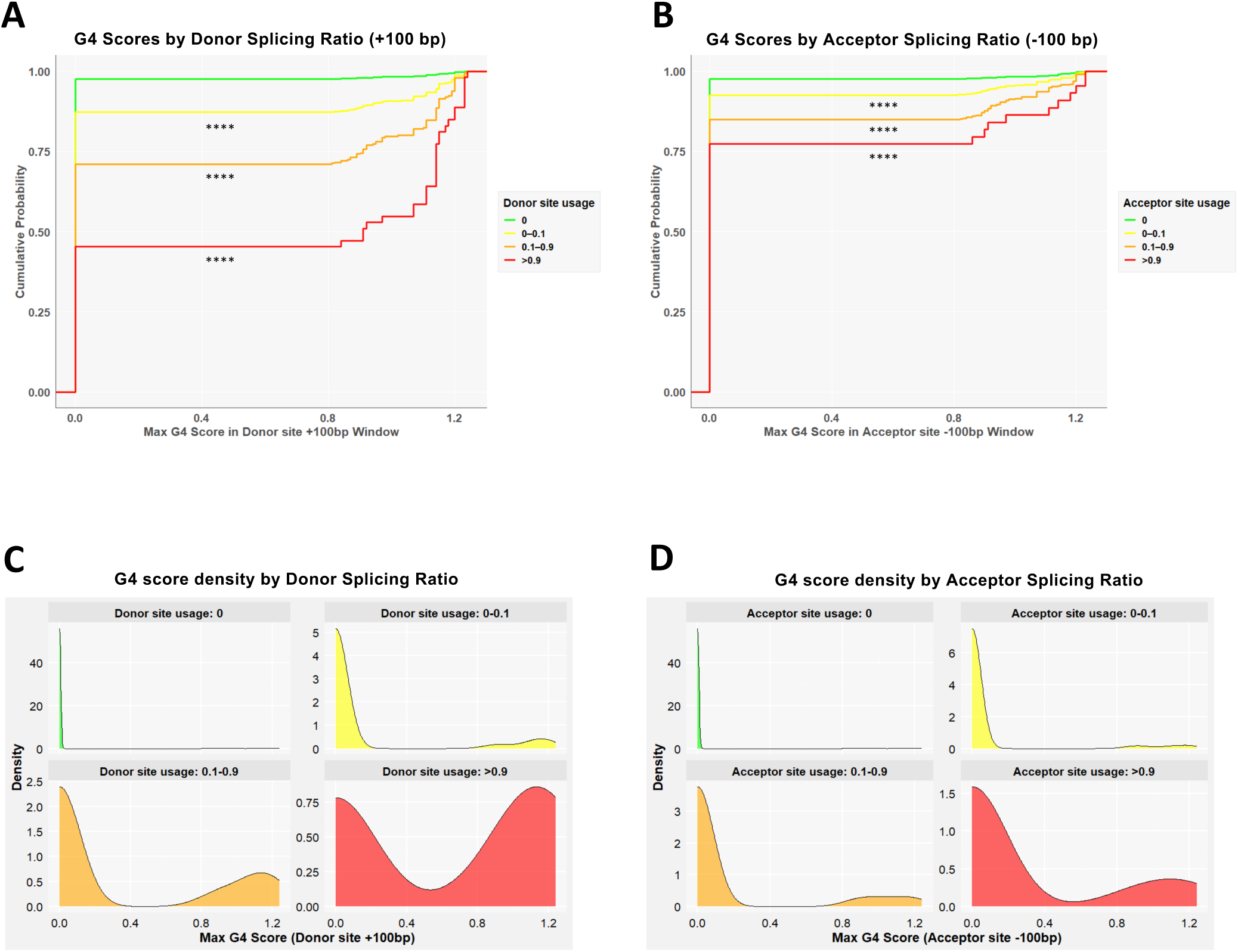
Correlation of G-quadruplex (G4) scores with splice site usage in the GFP_optimized library. To assess the influence of G-quadruplexes (G4) on splicing, we calculated G4Hunter scores at every nucleotide and recorded the maximum score within +100 nt downstream of each donor site or −100 nt upstream of each acceptor site. (**A, B**) Cumulative distribution plots show the maximum G4 scores stratified by splice site usage for donor (**A**) and acceptor (**B**) sites. Sites with high splicing activity (>0.9, red) show notable higher G4 scores for both donor and acceptor sites, revealing a modulatory role of G4 structures in splicing regulation. Significance: p < 0.0001 (****). (**C, D**) Density plots of the same G4 scores for donor (**C**) and acceptor (**D**) sites reveal distinct score distributions across splicing ratio groups. Curves shift rightward with increased splicing, suggesting higher G4 propensity at more frequently used splice sites.

**Supplementary Figure S13.**
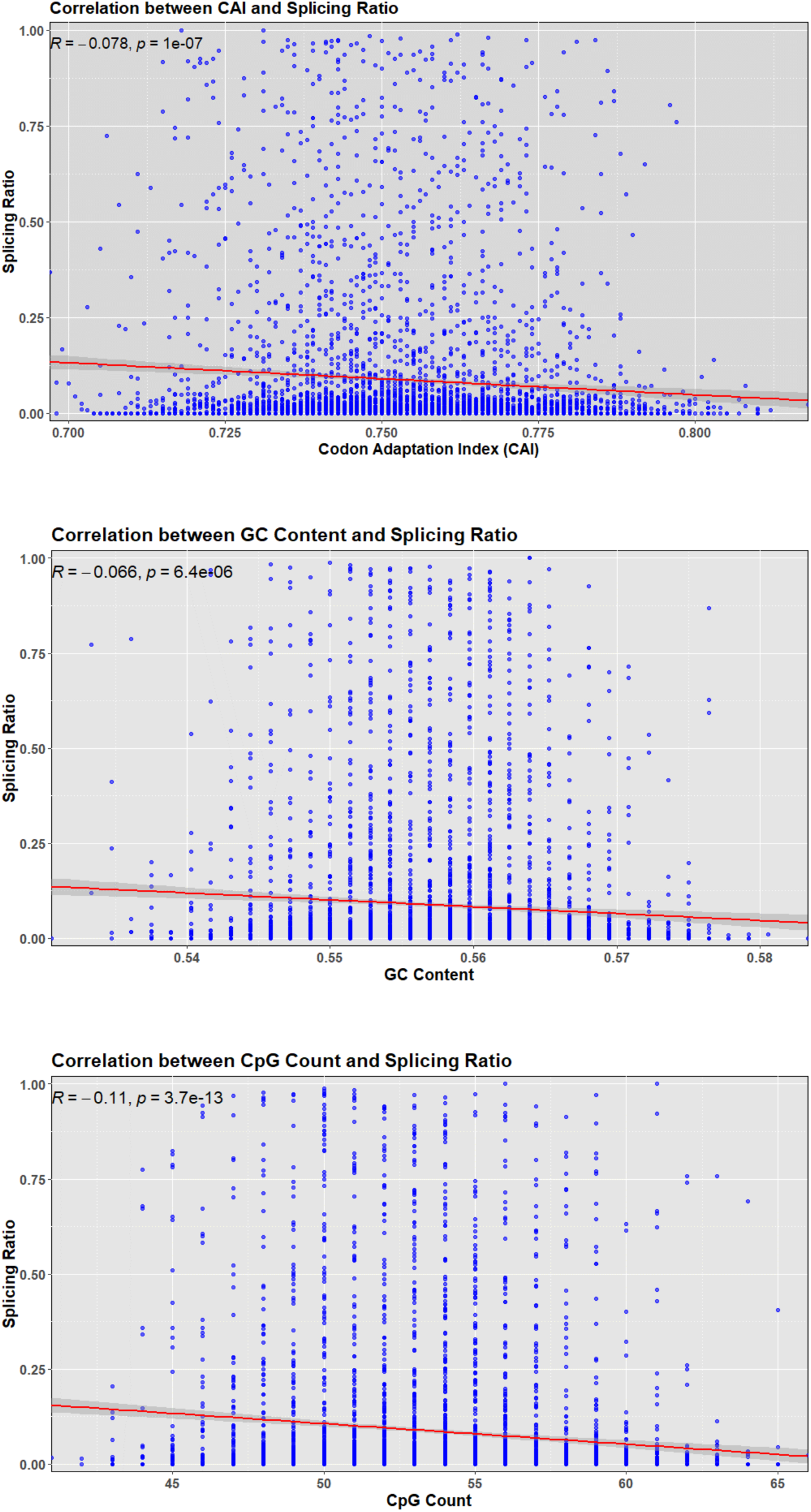
Correlation of Codon Adaptation Index (CAI), GC content and CpG count with splicing in the GFP_optimized library. Scatter plots show the relationship between the GFP mRNA splicing ratio and three sequence-derived features across all GFP variants: Codon Adaptation Index (CAI, top), GC content (middle), and CpG count (bottom). Each point corresponds to a single GFP variant. Linear regression (red line with 95% confidence interval in grey) reveals a weak but significant negative correlation between splicing and all three features: CAI (R = −0.078), GC content (R = −0.066), and CpG count (R = −0.11).

**Supplementary Figure S14.**
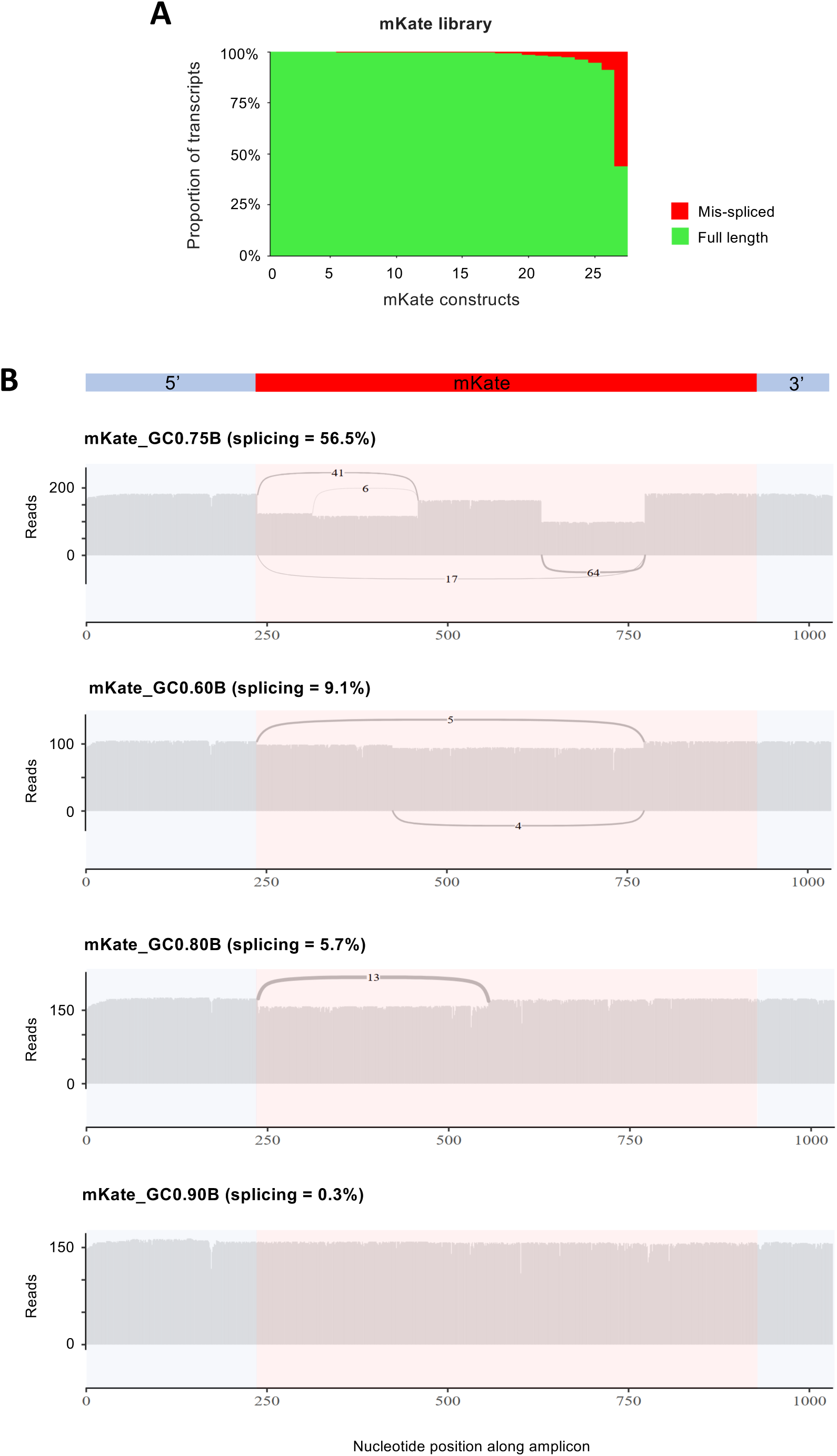
Splicing of synonymous mKate variants. **(A)** Stacked bars show the fraction of spliced (red) versus full-length (green) cDNA for each variant in the mKate library (n=27). **(B)** Sashimi plots for the three variants with the highest splicing (top) and one variant with residual splicing (bottom). Each sashimi plot contains ∼150 random reads from a single barcode per mKate variant.

**Figure S15.**
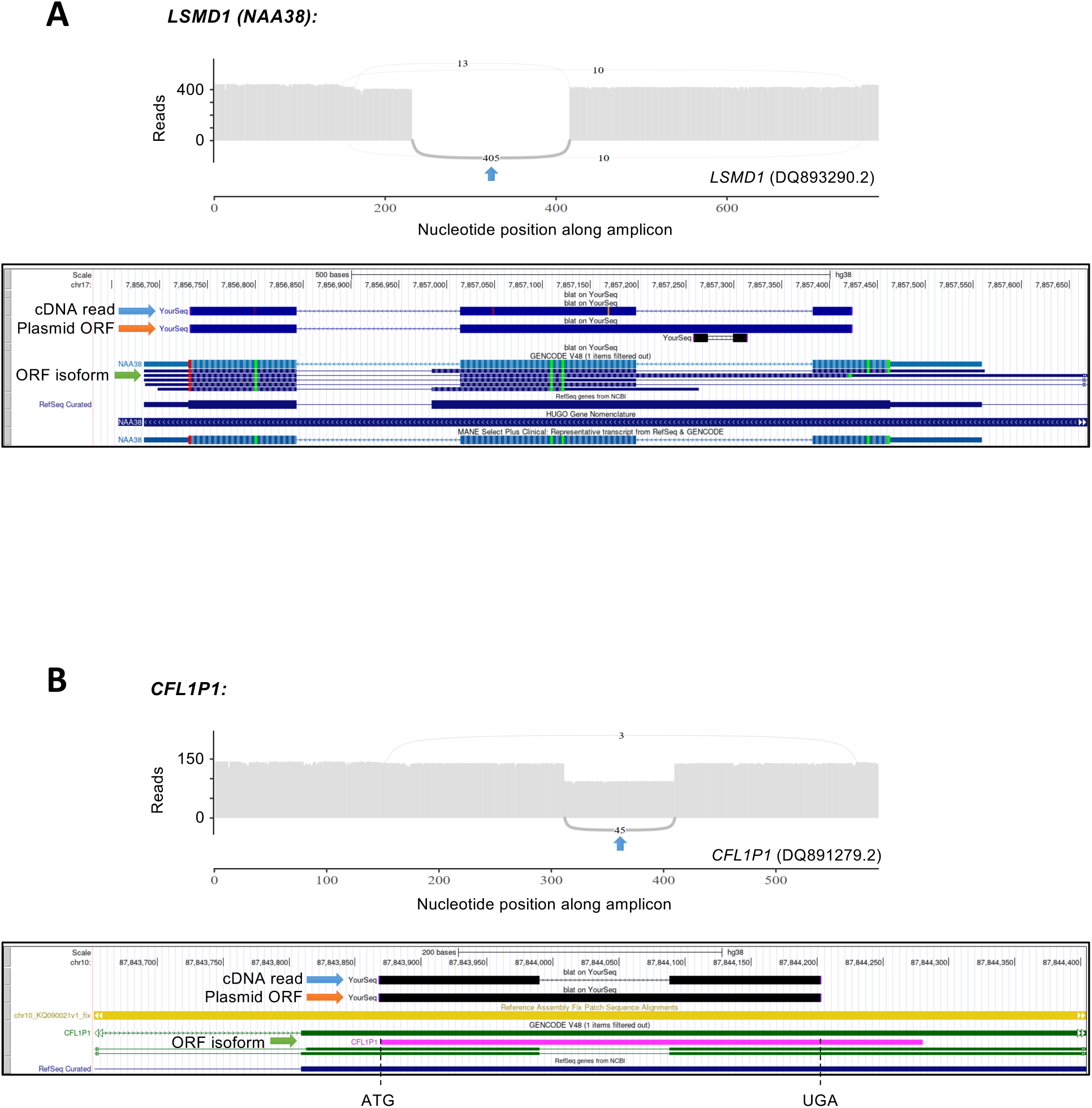
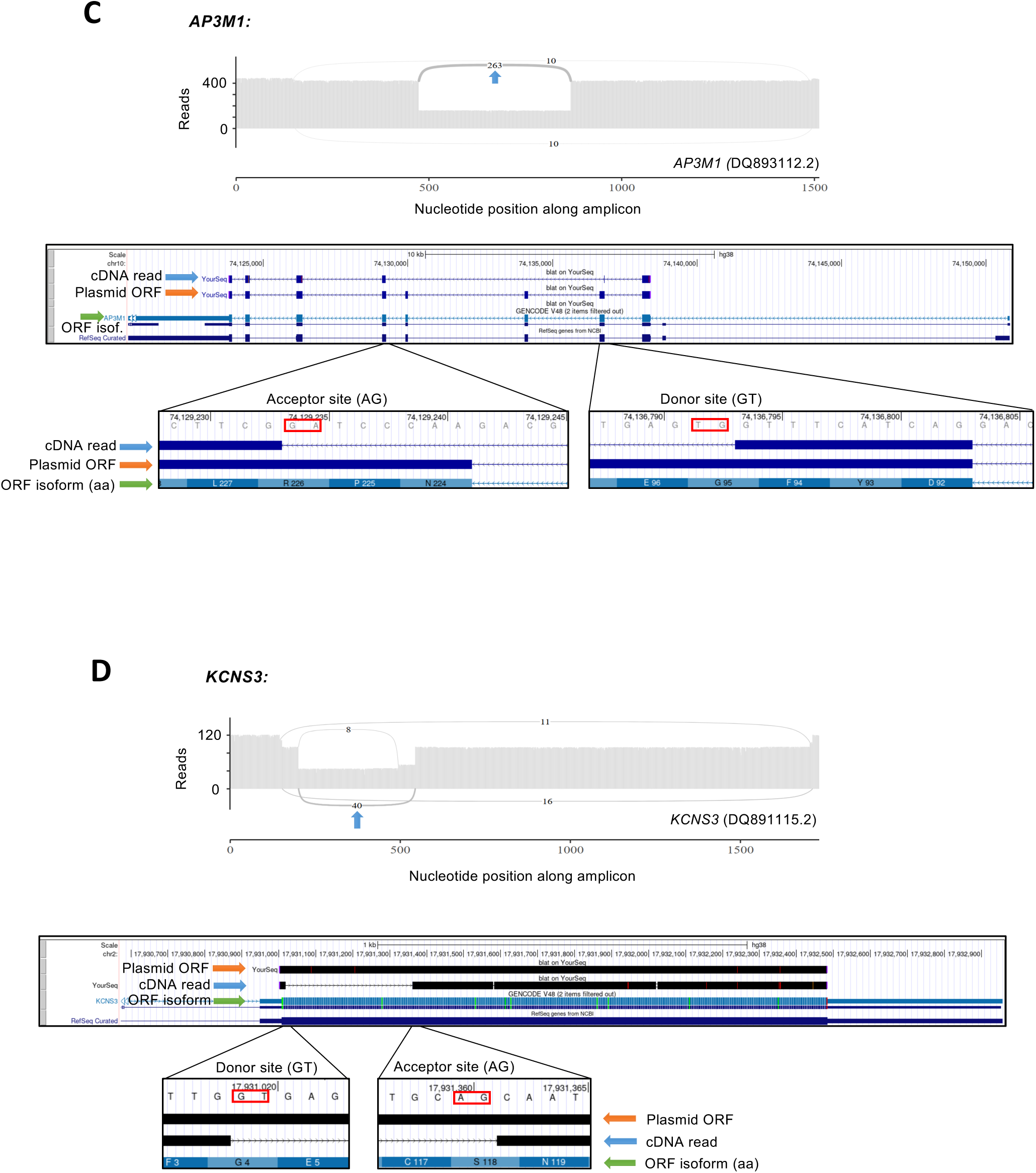
Representative splicing outcomes in ORFeome cDNAs. In each panel, the top track shows a sashimi plot where blue arrows point out the most common spliced isoform. Below, Genome Browser snapshots display BLAT alignments: blue arrows mark a representative read for the major spliced isoform; orange arrows denote the plasmid ORF reference sequence and green arrows indicate the endogenous reference isoform encompassed by the ORF insert. **(A,B)** Examples of ORF transcripts that recapitulated annotated splicing junctions: **(A)** DQ893290.2 – *LSMD1 (NAA38).* ORF corresponds to an *LSMD1* isoform with a retained intron relative to the principal *LSMD1* transcript. Long-read cDNA sequencing shows that nearly all transcripts splice across this region, recreating the annotated junction of the main isoform and yielding a hybrid full-length transcript. **(B)** DQ891279.2 – *CFL1P1.* The vector carries an ORF for a *CFL1P1* isoform (Met→UGA ORF; currently recategorized as non-coding). A substantial fraction of cDNA reads recapitulate annotated junctions from two *CFL1P1* isoforms, thereby generating a spliced isoform from the intronless plasmid sequence. **(C, D)** Examples of ORF transcripts spliced upon the activation of cryptic splicing sites. **(C)** DQ893112.2 – *AP3M1.* The majority of spliced reads show exon 2→exon 5 skipping, consistent with activation of exonic cryptic sites. **(D)** DQ891115.2 - *KCNS3.* The detection of spliced transcripts from an intronless ORF underscores that recombinant DNA manipulation, through alterations in sequence context following ORF insertion into an expression vector, can be sufficient to disrupt the splicing regulatory balance and induce aberrant splicing.

**Supplementary Figure S16.**
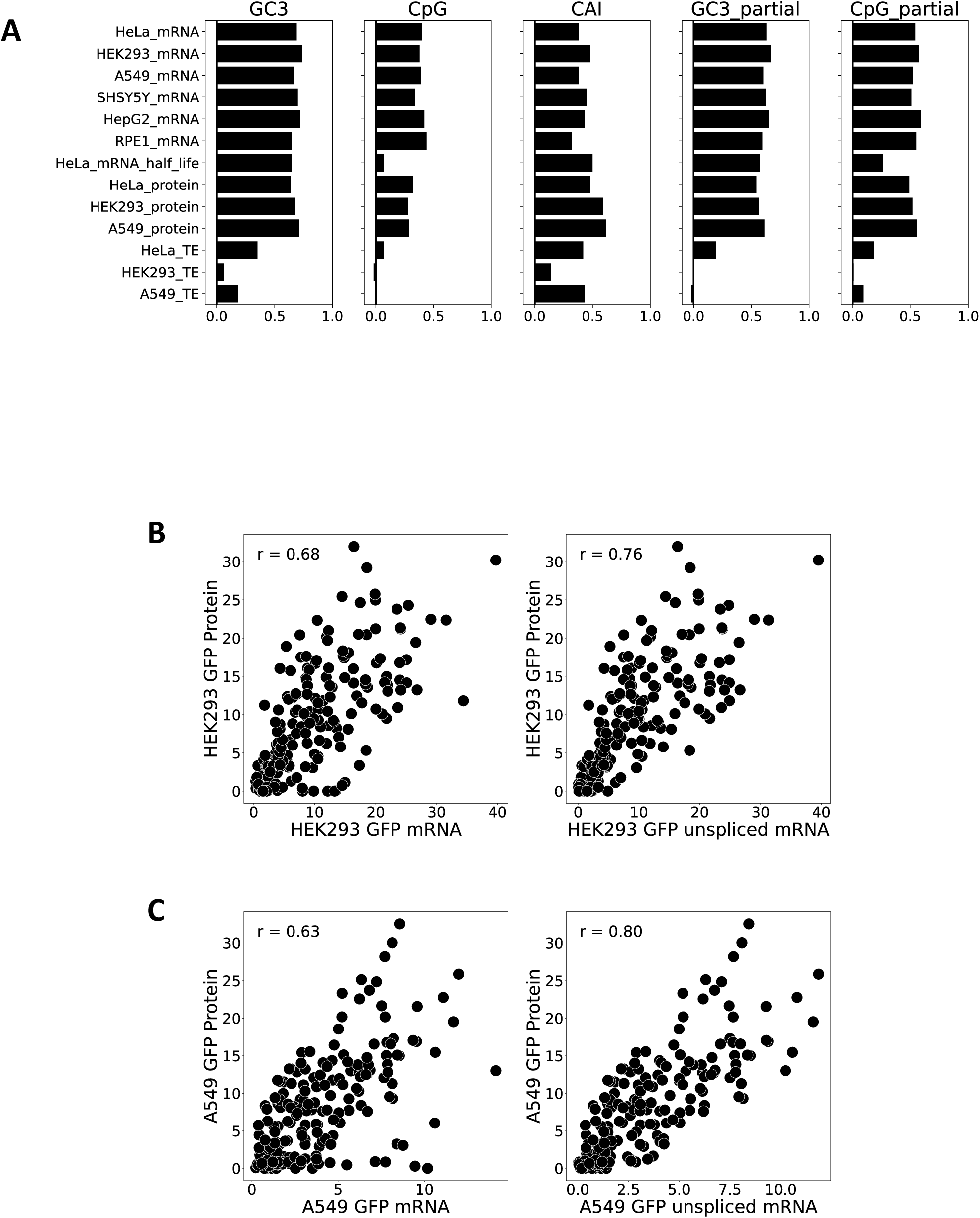
Accounting for splicing increases the correlation between mRNA and protein levels in HEK293 and A549 cells. **A)** Pearson correlations and partial correlations (controlling for CAI) of GFP gene features with expression levels. The first three panels were copied from Fig. 4B. **B)** Scatter plot showing correlation between GFP protein and total GFP mRNA (left) or estimated levels of full-length GFP mRNA (right) in HEK293 cells. **C)** Scatter plot showing correlation between GFP protein and total GFP mRNA (left) or estimated levels of full-length GFP mRNA (right) in A549 cells.

**Supplementary Figure S17.**
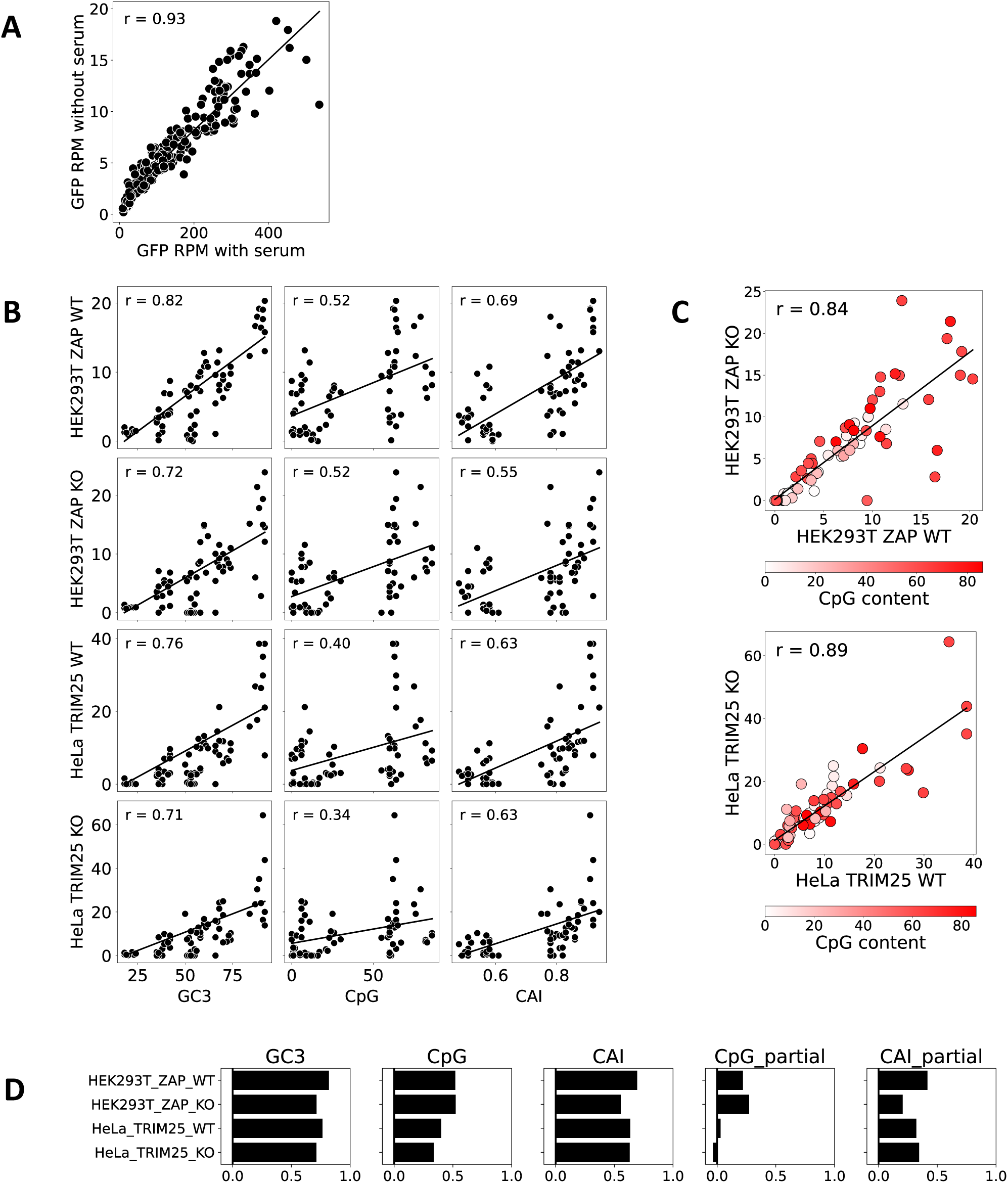
Consistency of expression patterns across treatment conditions and cells differing in ZAP pathway status. **A)** Correlation of GFP mRNA expression between serum-starved and serum-supplied HeLa cells. **B)** Correlations of the GFP protein expression level with gene features in HEK293 ZAP WT & KO cells, and HeLa TRIM25 WT & KO cells. **C)** Protein expression correlation between HEK293 ZAP WT & KO cells, and HeLa TRIM25 WT & KO cells**. D)** Pearson’s and Partial correlations of GFP gene features with expression levels in HEK293 ZAP WT & KO cells, and HeLa TRIM25 WT & KO cells. CpG_partial and CAI_partial were performed after controlling for the effects of GC3.

**Supplementary Figure S18:**
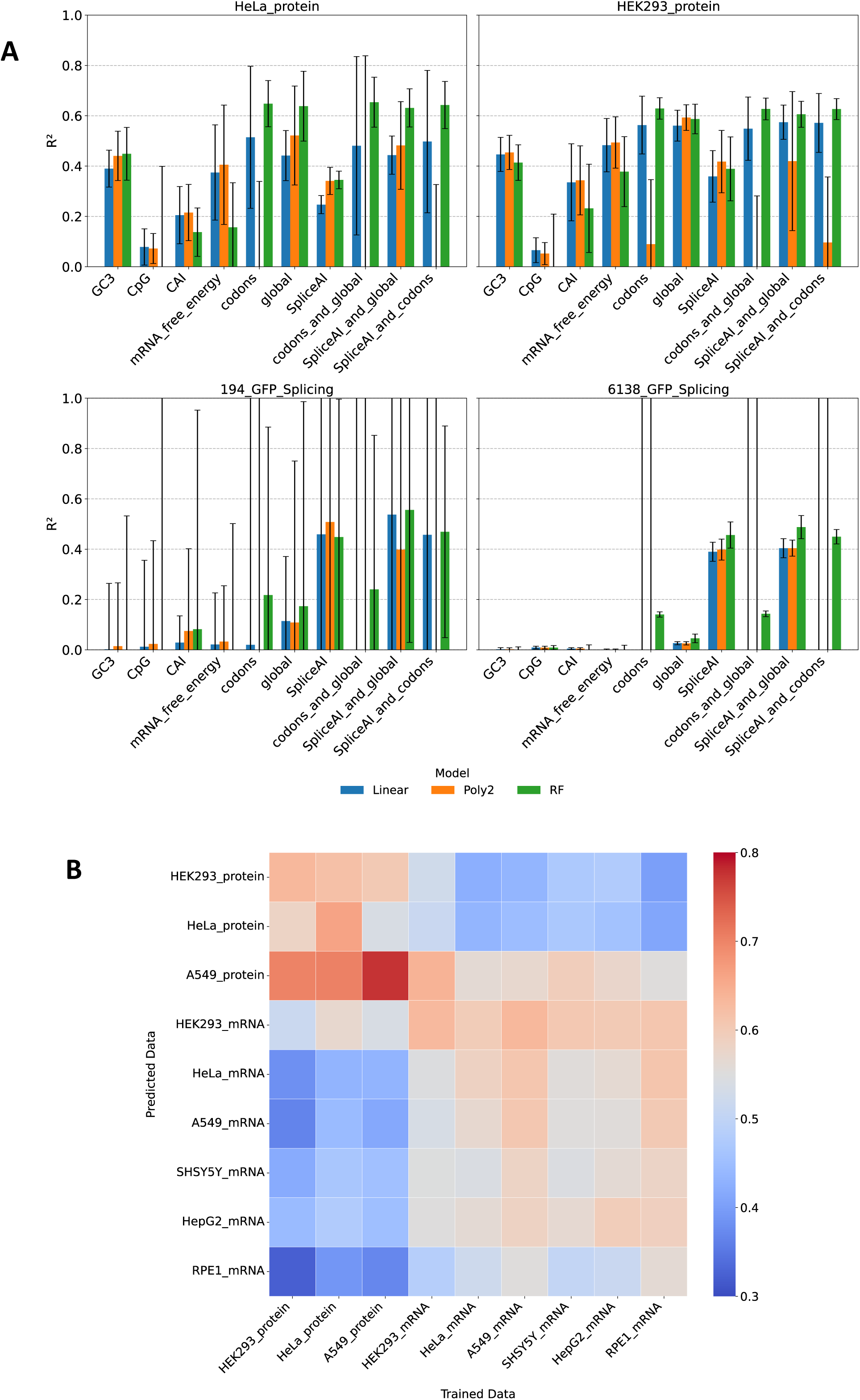
Predictive models of expression and splicing in HEK293 and HeLa cells. **A)** Performance on 20% unseen sequences of four model architectures in predicting GFP protein expression and splicing levels in A549 cells, using ten sets of predictive features. Top, prediction of protein levels, Bottom left, prediction of splicing in GFP_DOE library; bottom right, prediction of splicing in GFP_optimized library. **B)** Cross-modality predictions quantified as squared Pearson’s correlation coefficients across 9 sets of measurements and predictions. Out-of-fold predictions were generated using a random forest model trained on codons features with protein and mRNA expression data from multiple cell lines.

**Supplementary Figure S19:**
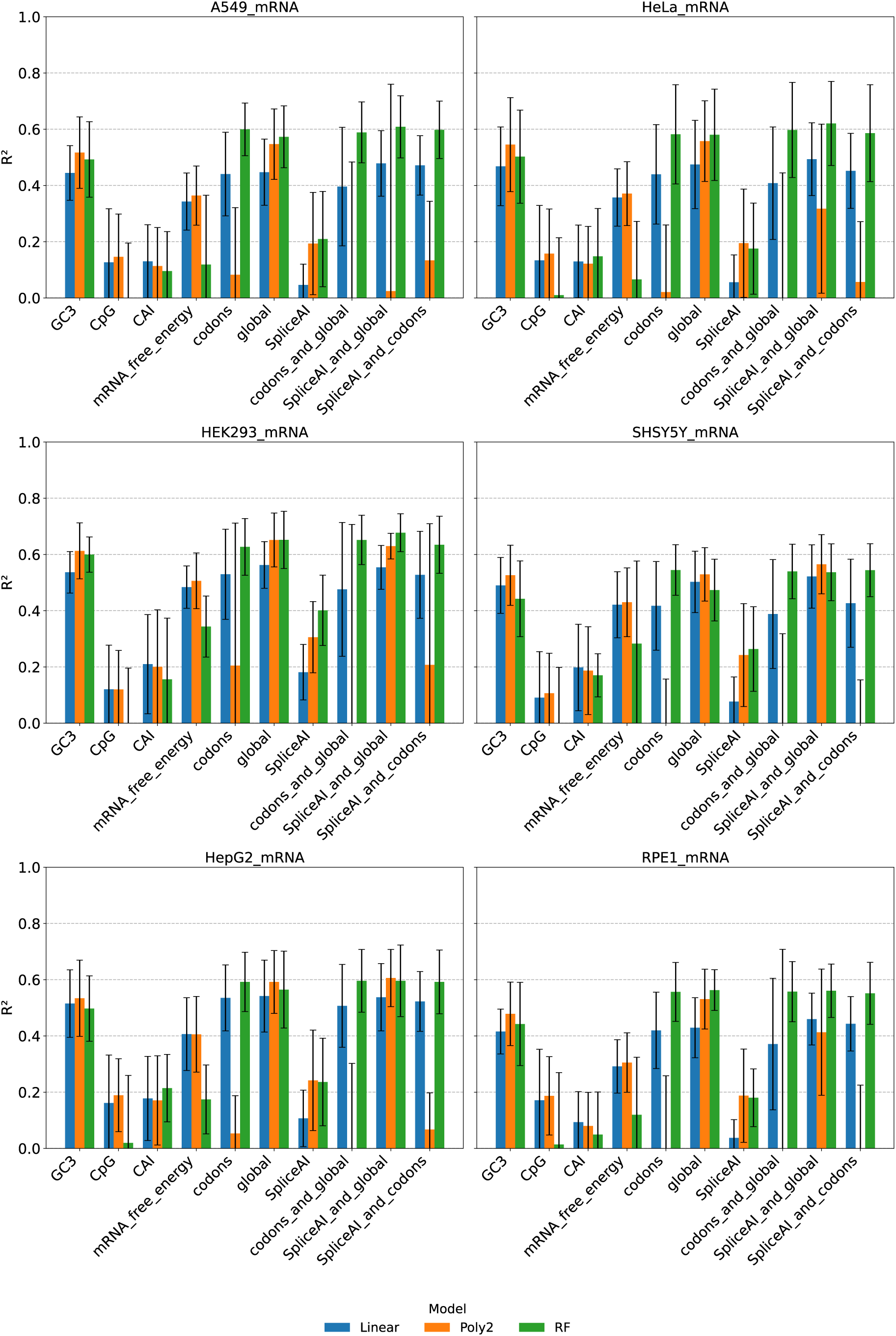
Predictive models of mRNA expression across cell lines.

**Supplementary Figure S20.**
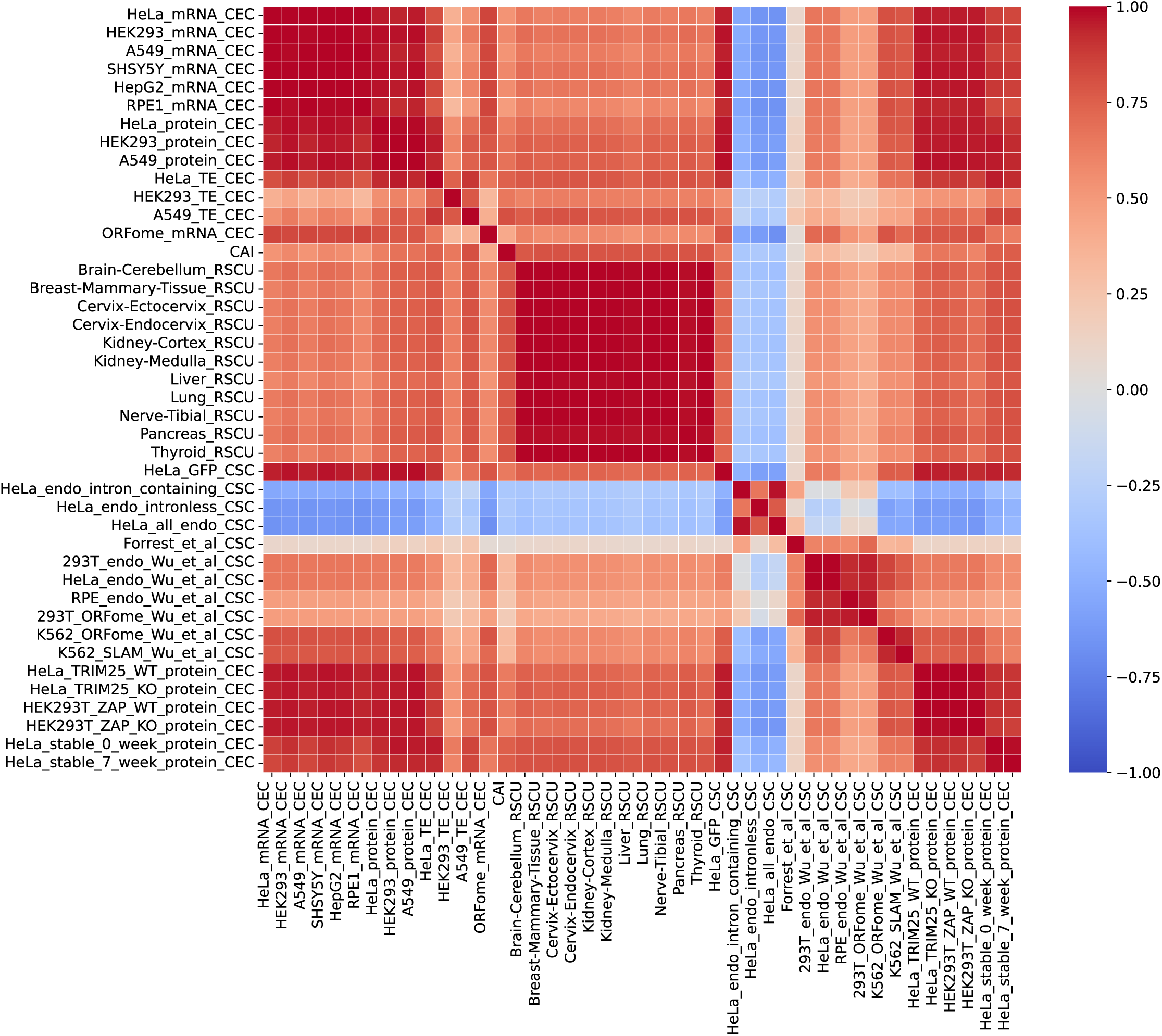
Correlation matrix of codon coefficients from this and previous studies.

---

**Table S1.**
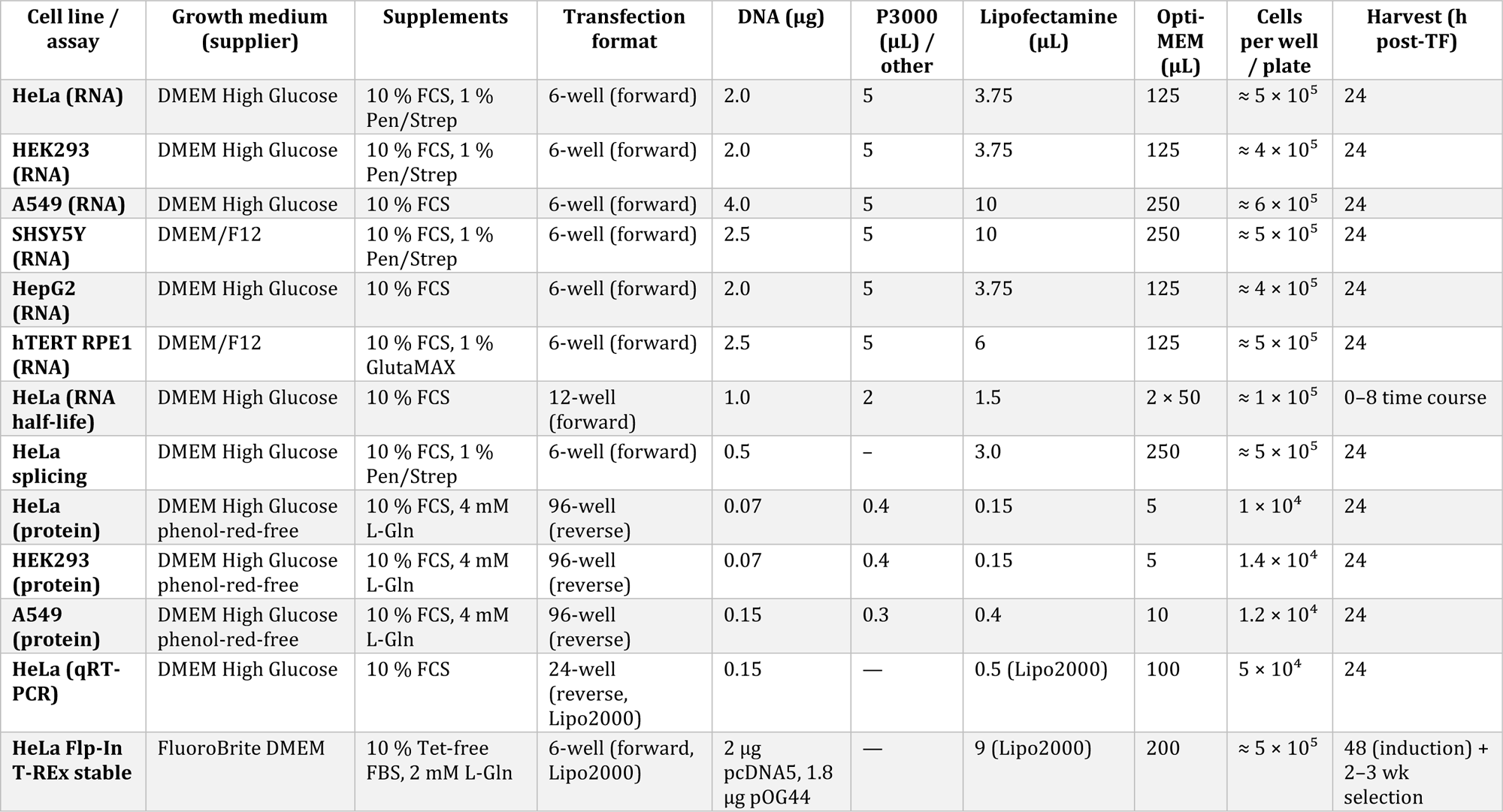
Cell culture and transfection conditions.

## Notes

### Competing Interest Statement

The authors have declared no competing interest.

